# A Virtual Nodule Environment (ViNE) for modelling the inter-kingdom metabolic integration during symbiotic nitrogen fixation

**DOI:** 10.1101/765271

**Authors:** George C diCenzo, Michelangelo Tesi, Thomas Pfau, Alessio Mengoni, Marco Fondi

## Abstract

Biological associations are often premised upon metabolic cross-talk between the organisms, with the N_2_-fixing endosymbiotic relationship between rhizobia and leguminous plants being a prime example. Here, we report the *in silico* reconstruction of a metabolic network of a *Medicago truncatula* plant nodulated by the bacterium *Sinorhizobium meliloti*. The nodule tissue of the model contains five spatially distinct developmental zones and encompasses the metabolism of both the plant and the bacterium. Flux balance analysis (FBA) suggested that the majority of the metabolic costs associated with symbiotic nitrogen fixation are directly related to supporting nitrogenase activity, while a minority is related to the formation and maintenance of nodule and bacteroid tissue. Interestingly, FBA simulations suggested there was a non-linear relationship between the rate of N_2_-fixation per gram of nodule and the rate of plant growth; increasing the N_2_-fixation efficiency was associated with diminishing returns in terms of plant growth. Evaluating the metabolic exchange between the symbiotic partners provided support for: i) differentiating bacteroids having access to sugars (e.g., sucrose) as a major carbon source, ii) ammonium being the major nitrogen export product of N_2_-fixing bacteria, and iii) N_2_-fixation being dependent on the transfer of protons from the plant cytoplasm to the bacteria through acidification of the peribacteroid space. Our simulations further suggested that the use of C_4_-dicarboxylates by N_2_-fixing bacteroids may be, in part, a consequence of the low concentration of free oxygen in the nodule limiting the activity of the plant mitochondria. These results demonstrate the power of this integrated model to advance our understanding of the functioning of legume nodules, and its potential for hypothesis generation to guide experimental studies and engineering of symbiotic nitrogen fixation.

## INTRODUCTION

Macroorganisms are colonized by a staggering diversity of microorganisms, collectively referred to as a ‘holobiont’ (1, 2). The intimate association between organisms is often driven by metabolic exchanges: many insects obtain essential nutrients from obligate bacterial symbionts (3), most plants can obtain phosphorus from arbuscular mycorrhiza in exchange for carbon (4), and the gut microbiota is thought to contribute to animal nutrition (5, 6). Complex global patterns often emerge during these intimate biological associations (7), especially when nutritional inter-dependencies are involved (8–10). The communication between the two metabolic networks of the interacting organisms may give rise to unpredicted phenotypic traits and unexpected emergent properties. Metabolic relationships can span over a large taxonomic range and have profound biological relevance (11–14). For example, the interactions between bacteria and multicellular organisms have been suggested to be key drivers of evolutionary transitions, leading to eukaryotic diversification and to the occupancy of novel niches (9, 15, 16). The study of the association of two biological entities is mainly challenged by the size of the system and by the unpredictability of their metabolic interactions. Theoretical, systems-level models are required to unravel the intimate functioning of metabolic associations and eventually exploit their potential in biotechnological applications.

Symbiotic nitrogen fixation (SNF) is a paradigmatic example of the importance and the complexity of natural biological associations. SNF is a mutualistic relationship between a group of plant families, including the Fabaceae, and a polyphyletic group of alpha- and beta-proteobacteria known as rhizobia, or a taxa of Actinobacteria (*Frankia* spp.), in which the plants provide a niche and carbon to the bacteria in exchange for fixed nitrogen (17). SNF involves constant metabolic cross-talk between the plant and the bacteria (18), and it is a paradigmatic example of bacterial cellular differentiation (19) and sociomicrobiological interactions (20). The rhizobia intra-cellularly colonize plant cells of a specialized organ known as a root (or stem) nodule. The intra-cellular rhizobia (referred to as bacteroids) are surrounded by a plant derived membrane, and the term symbiosome is used in reference to the structure consisting of the bacteroid, the plant derived membrane (i.e., the peribacteroid membrane), and the intervening space (i.e., the peribacteroid space). Nodules with an indeterminate structure, such as those formed by the plant *Medicago truncatula*, are divided into spatially distinct developmental zones (21) with a distal apical meristem and a proximal nitrogen fixation zone.

SNF plays a key role in the global nitrogen cycle and is central to sustainable agricultural practices by reducing the usage of synthetic nitrogen fertilizers whose application results in a multitude of adverse environmental consequences (22–24). Unfortunately, our ability to maximize the benefit of SNF is limited since rhizobial inoculants are often poorly effective due to low competitiveness (25, 26) and because rhizobium symbioses are specific to leguminous plants. Manipulating the rhizobium – legume interaction for biotechnological purposes will require an understanding of what we know and what we don’t know, as well as an ability to predict the consequences of genetic changes and environmental perturbations.

From a metabolic perspective, genome-scale metabolic reconstruction (GENREs) and constraint-based modelling has great potential to fulfill these roles. A GENRE also serves as a comprehensive knowledgebase of an organism’s metabolism, containing hundreds to thousands of metabolic and transport reactions, most of which are linked to the corresponding gene(s) whose gene product(s) catalyzes the reaction (27, 28). With the aid of mathematical approaches such as flux balance analysis (FBA), GENREs can be used to identify emergent system-level properties, to predict active reactions, and to identify essential genes (29). Compared to simple enrichment analyses that are typical in -omics studies, GENRE-based methods allow for the interpretation of data in a connected manner based on network topology and to infer the effects of changes in remote pathways on the overall cell physiology. When considering interacting entities, for example, this approach can predict the consequence of mutations in one organism on the metabolism of the other. However, multi-organism metabolic reconstructions are still in their infancy, and very few examples of combined models exist compared to single strain GENREs (8, 14, 30–32).

Despite the importance of metabolism to SNF (18), there has been limited use of metabolic modelling in the study of rhizobia and SNF. To date, GENREs of varying quality have been reported for only three rhizobia: *Sinorhizobium meliloti* (33–35), *Rhizobium etli* (36–38), and *Bradyrhizobium diazoefficiens* (39). Currently, *M. truncatula* (40) and *Glycine max* (41) are the only legumes with published GENREs. With the exception of the *G. max* GENRE, these GENREs have been used in preliminary analyses of SNF, providing results generally consistent with expectations. However, all analyses to date suffer from two major limitations. Simulations with the rhizobium models ignore plant metabolism, while simulations with the *M. truncatula* GENRE involved a very limited draft *S. meliloti* metabolic reconstruction. Furthermore, all simulations have focused on the final stage of SNF and have not considered the different steps of the preceding developmental progression where metabolism remains poorly understood (18).

Here, we report a holistic *in silico* representation of the integrated metabolism of the holobiont consisting of a *M. truncatula* plant nodulated by *S. meliloti*, which we refer to as a <u>Vi</u>rtual <u>N</u>odule <u>E</u>nvironment (ViNE). Our combined, multi-compartment reconstruction accounts for the metabolic activity of shoot and root tissues together with a nodule consisting of five developmental zones. We report initial characterizations of ViNE using FBA, including zone-specific metabolic properties, trade-offs between nitrogen-fixation and plant growth, and the usage of dicarboxylates as a carbon source by bacteroids. Going forward, we expect ViNE will provide a powerful platform for hypothesis generation aimed at understanding and quantitatively evaluating SNF, as well as guiding attempts at engineering SNF for increased symbiotic efficiency.

## MATERIALS AND METHODS

### Preparing an improved *S. meliloti* metabolic reconstruction

A new *S. meliloti* metabolic reconstruction was built using the existing core metabolic reconstruction iGD726 (34) as a starting point. First, the biomass composition was updated as summarized in Table S1. In particular, glycogen was reduced to 0.1% cell dry weight (CDW), poly-hydroxybutyrate was reduced to 1% CDW, and high and low molecular weight succinoglycan were reduced to 0.1% and 0.4% CDW, respectively (42). Additionally, putrescine and spermidine were added to the biomass composition at trace concentrations (43).

The working reconstruction was manually expanded to contain accessory metabolic pathways following our previously reported workflow (34). Briefly, the reconstruction was expanded by the addition of one pathway at a time. For each pathway, all reactions were individually added to the model, the gene associations and reaction equations checked against literature sources, and where possible, each reaction was referenced (see File S2). Reactions were predominately taken from the previously published *S. meliloti* genome-scale metabolic reconstruction iGD1575 (33) when possible; otherwise, they were taken from the Kyoto Encyclopedia of Genes and Genomes (44), MetaCyc (45), ModelSEED (46), or MetaNetX (47) databases.

An automated expansion of the metabolic network was then performed. Using the ‘tncore_expand’ function of the Tn-Core Toolbox, all reactions absent in the working reconstruction but present in the *S. meliloti* genome-scale metabolic reconstruction iGD1575b (34) were transferred to the working reconstruction. Then, i) all unnecessary ‘source’ reactions were removed, ii) most metabolic reactions associated with an unknown gene were removed, and iii) some newly added reactions likely to be incorrect based on published literature were deleted. Reactions added during the automated expansion sharing a gene in common with an existing reaction were then manually examined, and in most cases manually removed from the reconstruction. Then, all reactions producing dead-end metabolites were iteratively removed,

The working reconstruction was next mass and charge balanced. Metabolite formulas and charges were obtained from the MetaNetX database (47) when available; otherwise, metabolite charges and formulas were manually prepared, using information from the PubChem database (48) when available. The ‘checkMassChargeBalance’ function of the COBRA Toolbox was used to identify mass or charge unbalanced reactions, and reaction equations were manually balanced. Duplicate reactions were identified and removed.

An ATP hydrolysis reaction was added to account for non-growth associated maintenance (NGAM) costs (49), using a NGAM cost of 8.39 mmol ATP h^−1^ (g dry weight)^−1^ as reported for *Escherichia coli* (50). A growth associated maintenance (GAM) reaction was not added as the reconstruction includes transcription and translation reactions. The final reconstruction, termed iGD1348, contains 1348 genes, 1407 reactions (1164 associated with at least one gene), and 1160 metabolites (Table S2). The final reconstruction is available in File S2 in SBML, XLS, and MATLAB COBRA format.

### Updating the *M. truncatula* metabolic network reconstruction

The published *M. truncatula* metabolic network reconstruction (40) was built based on the *M. truncatula* genome version Mt3.5v5 (51). Here, the gene associations were updated to correspond to the annotations of version 5.0, the most recent version of the *M. truncatula* genome (52). A conversion table was prepared linking the Mt3.5v5 gene names with the corresponding gene names from the Mt4.0v1 genome annotation (53), which were in turn associated with the corresponding gene names from the version 5.0 annotation. This conversion table was prepared based on the information present in i) the ‘Mt3.5-Mt4.0v1_conversion_table.txt’ file available on medicagogenome.org (54), and ii) the ‘MtrunA17r5.0-ANR-EGN-r1.6.gene-repeat_region.vs. JCVI-Mt4.0-gene.kgb.synonymy.txt’ file available online at medicago.toulouse.inra.fr/ MtrunA17r5.0-ANR. Next, the published *M. truncatula* SBML model was imported into MATLAB with the ‘importMedicago’ function of Pfau *et al*. (40). After importing, for genes with a one-to-one match between genome versions, the existing gene name was replaced with the gene name in the version 5.0 genome annotation. When multiple genes were combined into a single gene in the version 5.0 annotation, all of the genes were removed from the model and replaced with the single gene. Genes that were split into multiple genes in the version 5.0 annotation were replaced with all of the new genes using an ‘or’ association. Genes with no match in the Mt5.0 genome were removed from the model; reactions constrained upon removing these genes were also deleted unless they were essential for model growth (i.e., RXN-9944_H, RXN-7674_H, and PASTOQUINOL--PLASTOCYANIN-REDUCTASE-RXN_H), in which case the corresponding gene also was not removed.

The majority of transport reactions in the original *M. truncatula* metabolic reconstruction, both between the cell and the external environment and between organelles, were simple diffusion reactions lacking an energy source such as ATP hydrolysis or proton cotransport. To limit inappropriate transport between compartments, all single-metabolite diffusion reactions were modified with the exception of metabolites such as water, gases, and light. All bidirectional reactions were split into two unidirectional reactions, and each reaction was modified to require the hydrolysis of 0.25 mol of ATP per mol of transported compound. The modified reconstruction contains 2522 genes, 2920 reactions (1722 associated with at least one gene), and 2742 metabolites.

The updated *M. truncatula* reconstruction was used to generate a tissue-specific *M. truncatula* model containing shoot and root tissues using the ‘BuildTissueModel’ function of Pfau *et al.* (40). Reactions to transfer metabolites between the root and shoot tissue were modified to require the hydrolysis of 0.25 mol of root ATP and 0.25 mol of shoot ATP per mol of transferred metabolite. The model was then modified to contain unique gene names for those associated with the shoot tissue and for those associated with the root tissue, following which all unused genes were removed from the model. Finally, root import reactions for the following compounds were added in anticipation of integration with the *S. meliloti* model: Co^2+^, MoO_4_^3^−, Mn^2+^, Zn^2+^, Ca^2+^, K^+^, and Na^+^. The final model encompassed root metabolism and shoot metabolism with appropriate cross-talk between the tissues (40), and all reactions, metabolites, and genes associated with the shoot contain the prefix ‘Leave_’, while those associated with the root contain the prefix ‘Root_’.

### Reconstructing the metabolism of a nodulated *M. truncatula* plant

The original full (i.e., non-tissue-specific) *M. truncatula* reconstruction (40) was imported to MATLAB in COBRA format from SBML format using the ‘readCbModel’ function. The model was updated to the version 5.0 genome annotations as described in the previously section, and diffusion reactions were modified to require an energy source as described in the previous section. The following reactions were then added in preparation for integration with the *S. meliloti* model: a homocitrate synthase reaction, a biotin source reaction, a H_2_ export reaction, and import reactions for each of N_2_, Mn^2+^, Zn^2+^, Ca^2+^, K^+^, and Na^+^. The gene *MtrunA17Chr1g0213481* was associated with the homocitrate synthase reaction based on homology to the gene of *Lotus japonicus* (55). At the same time, the *S. meliloti* model was modified such that fluxes were recorded in µmol hr^−1^ (g dry weight)^−1^, with one µmol of biomass equalling one g of biomass. This was done to ensure consistency with the units in the *M. truncatula* model. The *S. meliloti* model contained a single gene for all unknown GPRs (i.e., ‘Unknown’) and a single gene for all spontaneous reactions (i.e., ‘Spontaneous’). In preparation for constraining the nodule, the ‘Unknown’ and ‘Spontaneous’ genes were replaced with a series of genes each associated with a single reaction.

The following strategy was adopted to build a multi-compartment metabolic model accounting for the metabolic interactions of the two organisms. First, we mapped the two reconstructions to the same name space using the MetaNetX version 3.1 source files (47). This step was necessary as the *S. meliloti* model is based on the SEED database (56) and the *M. truncatula* model on the MetaCyc database (45). To minimize the required adjustments, only the metabolite identifiers of metabolites that were both i) a boundary metabolite in the *S. meliloti* model and ii) a cytoplasmic compound in the *M. truncatula* model were changed to the corresponding MetaNetX code; these represent the pool of metabolites that can be exchanged between the organisms. A multi-compartment reconstruction was built that included the *M. truncatula* model, the *S. meliloti* model, and transport reactions (without gene associations) that convert the *M. truncatula* cytoplasmic compounds to extra-cellular *S. meliloti* compounds (e.g., H2O_C => cpd00001[e]). These reactions represent the transport of compounds across the peribacteroid membrane, between the *M. truncatula* cytoplasm and the peribacteroid space of the symbiosome. For each metabolite, two unidirectional transport reactions were added that each required the hydrolysis of 0.25 mol of plant ATP per mol of the metabolite of interest. The exception was ammonia; in this case, the transport reaction into the peribacteroid space was driven by the hydrolysis of 0.25 mol of ATP per mol of ammonia, while the transport from the peribacteroid space was driven by proton symport (one proton per one molecule of ammonia). Additionally, protons transferred to the peribacteroid space from the *M. truncatula* cytoplasm were separated from protons exported by *S. meliloti*; no exchange of protons between *S. meliloti* and *M. truncatula* was allowed at this stage.

Four copies of the integrated *M. truncatula* – *S. meliloti* model were prepared to represent four distinct developmental zones of the nodule: zone II distal, zone II proximal, interzone II-III, and the nitrogen-fixing zone III. Additionally, a version of the *M. truncatula* model prior to integration with *S. meliloti* was included to represent zone I (apical meristem). In each of the five models, prefixes were added to all reactions, metabolites, and genes to specify to which zone and which organism the feature belongs (e.g., ‘NoduleIII_’ and ‘BacteroidIII_’). Next, all *S. meliloti* exchange reactions and all *M. truncatula* transport reactions were deleted in each of the five models. The exception was for nodule zone III, where the import of N_2_ and export of H_2_ by *M. truncatula* were not removed. Finally, a single model was produced that joined the tissue-specific (root and shoot) *M. truncatula* model with the five nodule zone models as a single COBRA formatted metabolic model. To this model, an irreversible reaction converting protons in the peribacteroid space to *S. meliloti* periplasmic protons was added specifically in nodule zone III, thereby allowing the transfer of protons fro *M. truncatula* to *S. meliloti*.

At this point, it was necessary to metabolically connect the nodule to the root and to the external environment. First, for each compound that could be exported by the *M. truncatula* root tissue, a reaction was added to each of the five nodule zones for the export of that compound. Then, for all compounds that could be imported by the *M. truncatula* root tissue (except ammonium and nitrate), a diffusion reaction (without an energy requirement) was added for the import of the metabolite from the external environment to a general nodule compound. Next, all compounds were identified that could be transferred between the root and shoot tissues in either direction. For each of these compounds, a diffusion reaction (without an energy requirement) was added to convert the compound in the root to a general nodule compound. Then, for each of the general nodule compounds, five irreversible reactions were added to transfer the general nodule metabolite to each of the nodule zones; each reaction involved the hydrolysis of 0.25 mol of nodule zone ATP per mol of transported metabolite. Finally, reactions were added to individually transfer asparagine and glutamine from the *M. truncatula* plant cytoplasm of nodule zone III (the nitrogen-fixing zone) to the root tissue, with each reaction requiring the hydrolysis of 0.25 mol of root ATP and 0.25 mol of nodule zone III ATP per mol of metabolite.

A series of biomass reactions were added to the combined model. A zone-specific biomass reaction was added to each of zone II distal, zone II proximal, and interzone II-III by combining *M. truncatula* and *S. meliloti* biomasses at a 75 : 25 ratio. Biomass of zone I consisted of only *M. truncatula* biomass. No biomass reaction was added to zone III as the purpose of this zone was to fix nitrogen. Next, an overall nodule biomass reaction was prepared by combining zone I, zone II distal, zone II proximal, and interzone IZ biomass at a 5 : 45 : 45 : 5 ratio. A plant biomass reaction was also prepared by combining shoot and root biomass at a 66.7 : 33.3 ratio. Finally, an overall biomass reaction was prepared that combined plant biomass with nodule biomass at a 98 : 2 ratio. The overall biomass reaction was set as the objective function during all FBA simulations unless stated otherwise.

All reactions that produced dead-end metabolites were iteratively removed from the model, followed by the addition of several constraints into the model (the list of the reactions removed following this procedure are listed in Dataset S1). Maintenance costs, in the form of ATP hydrolysis, were added to each tissue including the nitrogen-fixing zone III. The maintenance cost value for the shoot and root tissues were set as described elsewhere (40). Maintenance costs for plant nodule tissues were based on the shoot plus root maintenance costs scaled by the percent of biomass that consisted of the given nodule zone. Similarly, the maintenance costs of the bacteroid nodule tissues were based on a maximum of 50.4 µmol hr^−1^ (g plant dry weight)^−1^, scaled according to the percent of biomass that consisted of the given bacteroid zone (this value was chosen as it equals 30% the commonly used value for free-living *Escherichia coli*). Import of ammonium and nitrate by the root and nodule tissues was turned off, as was usage of starch as a carbon source in the shoot tissue. The uptake of light was set to 1000 µmol hr^−1^ (g plant dry weight)^−1^, which is within the range where there is a linear relation between light and CO_2_ usage (not shown). The total rate of oxygen usage by the plant and bacterial cells of nodule zone III was limited to 8.985 µmol hr^−1^ (g plant dry weight)^−1^. This value was arrived at as follows: i) the total oxygen usage of the entire nodule was limited to 12.98 µmol hr^−1^ (g plant dry weight)^−1^ based on published experimental data (57), ii) plant growth was optimized, iii) the O_2_ usage of zone III was limited to the O_2_ uptake rate in the initial analysis, and iv) the constraint on whole nodule O_2_ usage was removed. To force the use of C_4_-dicarboxylates by the bacteroids of zone III, reactions for the import of all other carbon sources into the symbiosomes were deleted. No constraints were pre-set on the transfer of nutrients from the plant cytosol to the bacteria of zone II distal, zone II proximal, or interzone IZ. Finally, the upper and lower bounds of all reactions were multiplied by 1000, converting the units to nmol hr^−1^ (g dry weight)^−1^. This step was necessary to avoid numerical issues when running GIMME due to low fluxes through the bacteroid reactions.

The reaction space of each nodule zone was constrained based on the *M. truncatula* – *S. meliloti* zone-specific RNA-seq data of Roux and coworkers (58), reanalyzed as described below, to obtain transcript per million (TPM) values. The expression threshold for a gene to be considered highly expressed was determined separately for each species, and it was equal to 1.1 times the average TPM value across all nodule zones of all genes that had at least one mapped read in at least one zone. To limit artificial differences between zones due to the choice to threshold, Kruskal-Wallis tests, followed by post-hoc comparisons, were performed for each gene to determine statistically significant between-zone expression changes; this was performed using the ‘agricolae’ package in R (59). If i) the difference between two zones was not statistically significant, ii) only one of the two zone had an expression value above the expression threshold, and iii) the value in the second zone was at least 80% of the expression threshold, then the value of the second zone was modified to be above the expression threshold. Moreover, as we wished to only constrain the reaction space of the nodule zones, all shoot and root genes were given artificial values above the expression threshold in order to ensure they were considered highly expressed.

The combined model was constrained using a custom multi-species adaptation of the gene-centric TIGER (60) implementation of the GIMME algorithm (61), which is available in the Tn-Core Toolbox (23). In short, GIMME was modified to take multiple gene lists (one list per species), multiple TPM lists (one list per species), and multiple expression thresholds (one per species). Genes above the respective expression threshold were considered expressed, and those below the respective threshold were turned off. A score for each ‘off’ gene was calculated by subtracting the expression value of each gene from the appropriate threshold. The scores for the species were then normalized based on the ratio of the expression thresholds. The normalized values of both species were combined as a single list, and the GIMME algorithm continued as normal. The growth fraction threshold for GIMME was set to 0.99.

The GIMME output was used as the basis to build a constrained and functional COBRA-formatted model. As the genes identified as ‘on’ following the GIMME analysis were insufficient to rebuild a working COBRA model, the following pipeline was used. All reactions active during the GIMME analysis with an absolute flux > 1 × 10^−6^ nmol hr^−1^ (g dry weight)^−1^ were identified. FASTCORE (epsilon of 1.01 × 10^−6^) was then run using these reactions as the input core reaction set and the same model used as input for GIMME, but with the lower bound of the biomass reaction set to 99% of the objective value. A list of protected reactions was prepared by combining: i) the output reactions of FASTCORE, ii) all reactions that were not constrained when the genes identified as ‘off’ in the GIMME analysis were deleted from the input model, iii) all peribacteroid transport reactions, and iv) all reactions for the transfer of metabolites between tissues. All nodule or bacteroid reactions that were not part of this protected list were removed from the model, and all genes no longer associated with a reaction were deleted. The genes associated with each reaction were then refined based on the GIMME output. For any given reaction, no change was made if all the associated genes were classified as ‘on’, or if all genes were linked with ‘and’ statements. Otherwise, for reactions with ‘or’ statements, but lacking ‘and’ statements, all genes classified as ‘off’ were removed from the reaction; if no gene was classified as ‘on’, then all genes were deleted except for the gene with the highest expression value. For reactions with both ‘or’ and ‘and’ statements, a complex loop was prepared. Put briefly, a minimal set of genes required for the reaction to be functional was left associated with the reaction, favouring the inclusion of ‘on’ genes followed by the inclusion of highly expressed ‘off’ genes. All reactions producing dead-end metabolites were iteratively removed from the model, and all genes no longer associated with a reaction were deleted.

Finally, all reaction and metabolite identifiers were updated to MetaNetX codes, where possible, to maximize consistency throughout the model, and duplicate reactions were deleted. We refer to this final version of the integrated model as ViNE (for <u>Vi</u>rtual <u>N</u>odule <u>E</u>nvironment), and it is provided in File S3 as MATLAB COBRA and SBML formatted files. The unconstrained model of the nodulated plant is also provided in File S3.

### Adding sucrose metabolism to zone III bacteroids in ViNE

To perform simulations comparing the use of sucrose and C_4_-dicarboxylates as a carbon source for zone III bacteroids, ViNE was modified as follows. The pipeline for construction of ViNE as detailed above was rerun with a single change. Prior to running GIMME, reactions for the import of all carbon sources, except sucrose, into the symbiosomes were deleted. This is in contrast to the construction of the regular version of ViNE, when the reactions for the import of all carbon sources, except C_4_-dicarboxlates, into the symbiosomes were deleted. The resulting model was then combined with ViNE, producing in an enlarged version of ViNE supplemented with the necessary reactions to allow sucrose to serve as a carbon source for zone III bacteroids.

### Analysis of the RNA-sequencing data

The nodule zones in the integrated metabolic model were constrained based on previously published RNA-seq data (58); however, the data were first re-analyzed using the *M. truncatula* version 5 genome sequence and annotations. The raw sequencing reads (fastq format; SRA accession SRP028599) were downloaded from the European Nucleotide Archive database (62), and all files corresponding to the same replicate of the same zone were concatenated as a single file, keeping separate files for each mate pair. The *M. truncatula* A17 genome (version 5.0) and the *S. meliloti* Rm2011 genome were downloaded and combined as a single file. The combined genome was indexed using the bowtie2-build function with default settings (63). Sequencing reads were mapped to the genome with bowtie2 version 2.2.3 (63), treating reads as paired-ends and using 20 threads. Output SAM files were sorted by name with samtools sort version 1.3.1-39-ga9054c7 (64) using 20 threads. Reads per gene were counted using HTseq-count version 0.6.1p1 (65), with the default alignment score threshold of 10. Each HTseq-count output table was split into two tables: one for *M. truncatula* and one for *S. meliloti*. TPM values were calculated for each species-specific output file using a custom Perl script, based on the total gene length for the *S. meliloti* genes and the total exon length for the *M. truncatula* genes. Finally, zone-specific average TPM values based the three biological replicates of each zone were calculated. These values were used to constrain the reaction space of the nodule zones in the integrated metabolic model.

### Metabolic modelling procedures

Model integration, model manipulations, and FBA simulations were performed in MATLAB R2016b (mathworks.com) using the SBMLToolbox version 4.1.0 (66), libSBML version 5.13.0 (67), and scripts from the COBRA Toolbox commit 9b10fa1 (68), the TIGER Toolbox version 1.2.0-beta (60), FASTCORE version 1.0 (69), and the Tn-Core Toolbox version 2.2 (70). The iLOG CPLEX Studio 12.7.1 solver (ibm.com) was used for nearly all FBA simulations; the exception was for preparation of iGD1348, during which the Gurobi version 7.0.1 solver (gurobi.com) was used. The switch to CPLEX was prompted by numerical issues that were solved by switching solver. All custom scripts used in this study are available through a GitHub repository (github.com/diCenzo-GC/ViNE_Reconstruction).

Each gene found in multiple tissues or nodule zones was distinguished by a unique gene name to facilitate tissue-specific gene deletion analysis. When performing global single or double gene deletion analyses, all versions of the gene were simultaneously deleted followed by the removal of all constrained reactions. In contrast, zone- or tissue-specific gene deletion analyses involved deleting just the gene version specific to the zone or tissue of interest. Flux variability was performed with the requirement that flux through the objective function was at least 99% the optimal flux. The robustness analysis involved first identifying the approximate flux range for each reaction in which the plant growth rate was non-zero. Then, for each reaction, the flux rate of the reaction was set to various values within the previously identified flux range, and the objective value was maximized. For simulations in which the rate of nodulation could vary, nodule biomass was removed from the objective reaction and instead forced through a nodule biomass sink reaction at the appropriate rate; maintenance costs and oxygen availability were modified accordingly (see Text S1). For simulations comparing the effect of providing zone III bacteroids sucrose versus C_4_-dicarboxylates as the carbon source, a modified version of ViNE was prepared as described in the subsection “Adding sucrose metabolism to zone III bacteroids in ViNE”.

## RESULTS

### Validation of iGD1348, an updated *S. meliloti* metabolic reconstruction

Prior to constructing the integrated plant – bacterium metabolic model, an updated metabolic reconstruction of *S. meliloti* Rm1021 was prepared as described in the Materials and Methods. Briefly, the highly refined core metabolic network iGD726 (34) was combined with the comprehensive accessory metabolism of the genome-scale metabolic network iGD1575 (33). Most of the reactions were compared against the literature, referenced where possible, and mass and charge balanced. The updated model consists of 1348 genes (Table S2), and incorporates information from 240 literature sources (listed in the Excel file of File S1) that include published transposon-sequencing (Tn-seq) data (34) and Phenotype MicroArray data (33, 71, 72) for wild-type and mutant strains.

Several tests were performed to validate the quality of the newly prepared *S. meliloti* reconstruction. Flux balance analysis (FBA) was used to simulate growth using glucose or succinate as the sole source of carbon, with or without the inclusion of an NGAM reaction. Inclusion of an NGAM reaction resulted in a specific growth rate reduction of ~ 0.043 h^−1^ and 0.030 h^−1^ for growth with glucose and succinate, respectively. This result confirmed the absence of energy leaks in iGD1348 that would allow for spontaneous energy production.

Using FBA, the ability of *S. meliloti* to catabolize 163 carbon sources to support growth was predicted with the iGD1348 and iGD1575 models (Dataset S2). As previously reported (33), simulations with the iGD1575 model correctly predicted growth with 67 of the 85 (79%) substrates experimentally shown to support growth of *S. meliloti*. Nicely, simulations with the iGD1348 model correctly predicted growth with 76 of these 85 (89%) substrates, including all 67 that supported growth of the iGD1575 model. This result confirmed that iGD1348 incorporates the majority of the accessory metabolism of *S. meliloti*, and that it is a better representation of total cellular *S. meliloti* metabolism than the previous genome-scale model.

Context-specific core metabolic models were extracted from the iGD1348 and iGD1575 metabolic models through the integration of transposon-sequencing (Tn-seq) data (34) using Tn-Core (29). The accuracy of the resulting core metabolic models was determined through comparison with iGD726, a manually prepared core metabolic model of *S. meliloti* (34). As summarized in Figure 1, the iGD1348 core model displayed greater overlap with the iGD726 model than did the iGD1575 core model. In particular, of the essential genes in the iGD726 model, 96% were essential in the iGD1348 core model, whereas only 62% were essential in the iGD1575 core model (Figure 1B). This result confirmed that the newly prepared iGD1348 reconstruction better represents the core metabolic network of *S. meliloti* than does the iGD1575 reconstruction. Overall, these tests confirmed that iGD1348 is a high-quality representation of *S. meliloti* metabolism, and that it better represents both the core and accessory metabolic properties of *S. meliloti* strain Rm1021 than does the original iGD1575 model.

**Figure 1.**
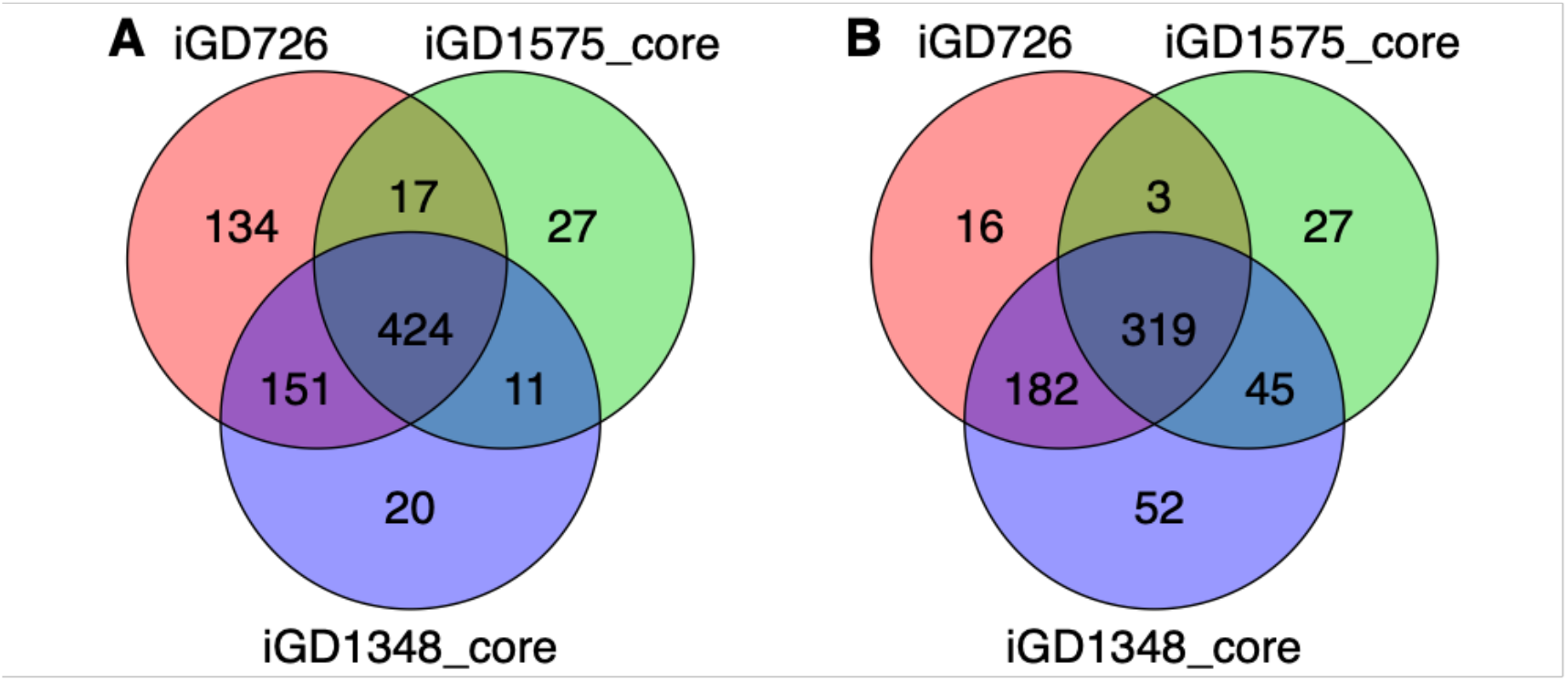
Overlap between the iGD726 model and core metabolic models derived from iGD1575 and iGD1348. Venn diagrams illustrating the overlap in (**A**) the total gene content, and (**B**) the essential genes of the following three models: the manually prepared iGD726 core model, a core model derived from iGD1575, and a core model derived from iGD1348. Core models of iGD1575 and iGD1348 were prepared using Tn-Core and published Tn-seq data(34).

### Construction and validation of a metabolic model of a nodulated legume

As a prerequisite to generating an *in silico* genome-scale metabolic network of an entire nodulated legume (referred to as ViNE for <u>Vi</u>rtual <u>N</u>odule <u>E</u>nvironment), it was necessary to obtain high-quality reconstructions of *M. truncatula* and *S. meliloti* metabolism. In the case of *M. truncatula*, we used a recently published reconstruction that was updated to match the most recent version of the *M. truncatula* genome annotation (see Materials and Methods). For *S. meliloti*, we made use of the newly updated model described in the previous section and the Materials and Methods.

Integrating the *S. meliloti* and *M. truncatula* metabolic models resulted in a model encompassing shoot, root, and nodule tissues as summarized in Figure 2 and Table 1. In total, this model includes 746 unique *S. meliloti* genes and 1,327 unique *M. truncatula* genes. Several simulations were performed to evaluate the reliability of the model. Using FBA, the maximal rate of plant (shoot + root) growth of the nodulated system was predicted to be ~ 0.044 g day^−1^ (g plant dry weight)^−1^, which is a reasonable prediction; *Medicago sativa* plants have an experimentally determined growth rate of ~ 0.1 g day^−1^ (g plant dry weight)^−1^ (73).

**Table 1.** Summary properties of ViNE.

| Model feature | Tissue |  |  |  |  |  |
| --- | --- | --- | --- | --- | --- | --- |
|  | Shoot | Root | Zone I | Zone II d | Zone II p | Zone IZ |
| **Genes** |  |  |  |  |  |  |
| *M. truncatula* | 1295 | 1292 | 236 | 265 | 243 | 228 |
| *S. meliloti* | 0 | 0 | 0 | 640 | 629 | 638 |
| **Reactions \*** |  |  |  |  |  |  |
| *M. truncatula* | 937 | 944 | 494 | 543 | 559 | 530 |
| *S. meliloti* | 0 | 0 | 0 | 662 | 654 | 670 |
| **Metabolites** |  |  |  |  |  |  |
| *M. truncatula* | 831 | 825 | 490 | 568 | 597 | 581 |
| *S. meliloti* | 0 | 0 | 0 | 751 | 747 | 764 |

\* Excludes reactions for transfer of metabolites between tissues or between *M. truncatula* and the peribacteroid space.

**Figure 2.**
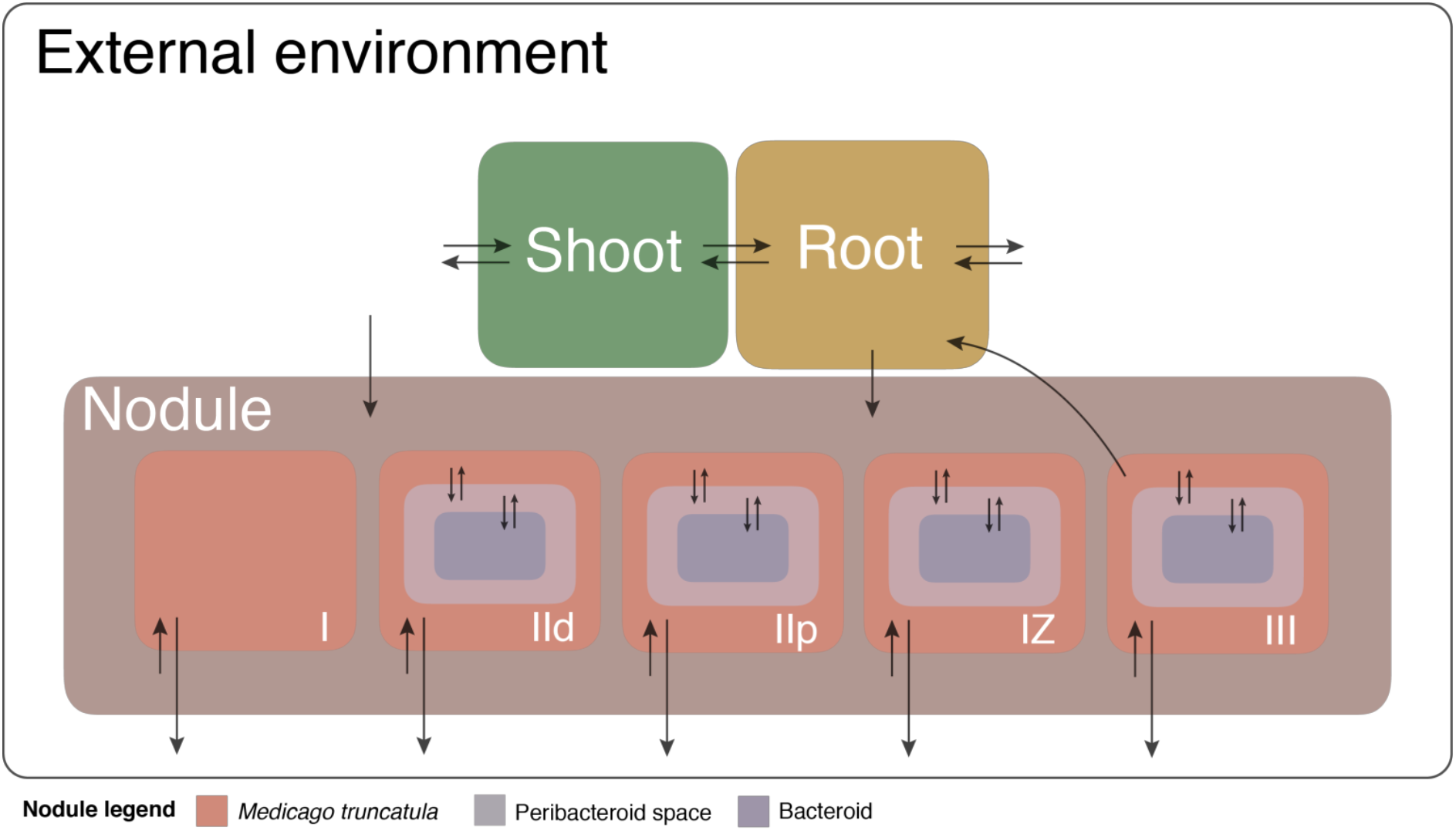
Visual depiction of ViNE. A schematic summarizing the overall structure of the *S. meliloti* nodulated *M. truncatula* plant developed in this work. The model contains three plant tissues (shoot, root, nodule) with the nodule subdivided into five developmental zones. Arrows indicate transport reactions with the direction representative of the directionality of the transport reactions. The scale of the figure has no meaning.

Next, FBA was used to examine the effects of adding exogenous ammonium to the soil on plant growth considering two situations: i) the rate of N_2_-fixation could vary while the rate of nodulation was constant, and ii) the rate of nodulation could vary while the rate of N_2_-fixation per gram of nodule was constant. As expected, increasing the availability of exogenous ammonium increased the rate of plant growth, with the effect more pronounced when nodulation was allowed to decrease with increasing ammonium since the plant no longer had to invest in nodule maintenance (Figure 3).

**Figure 3.**
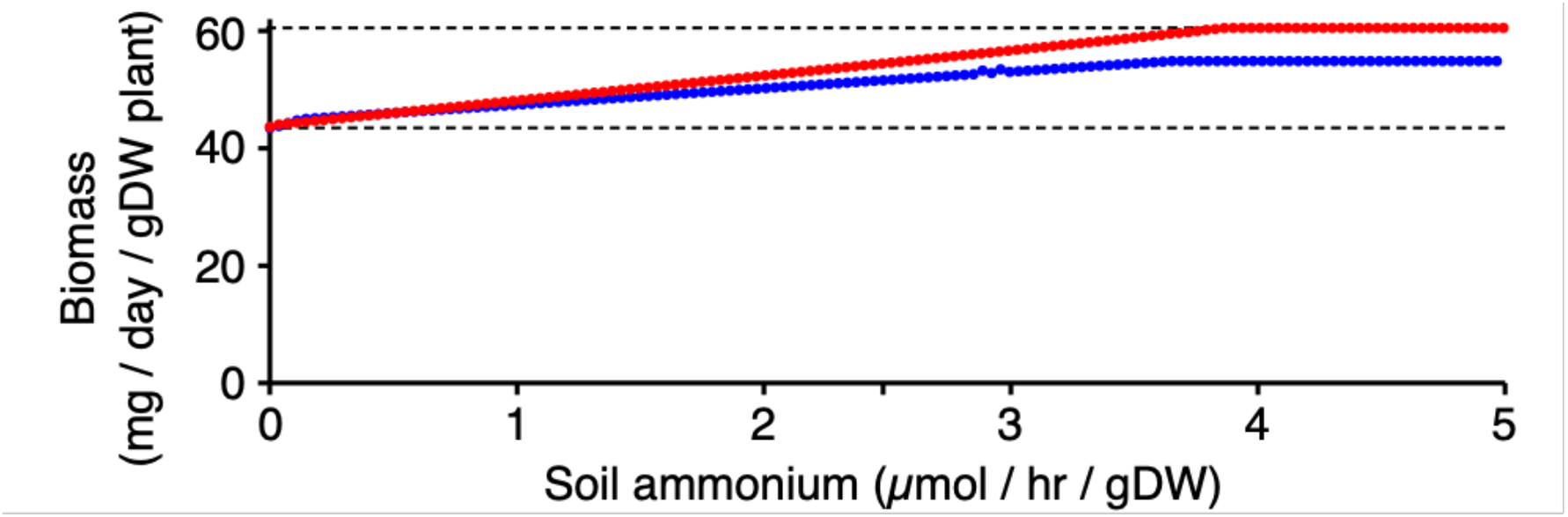
Effect of exogenous ammonium on *M. truncatula* growth. The effects of increasing the availability of soil ammonium on the growth rate of nodulated *M. truncatula* was examined. Simulations were run allowing either the rate of N_2_-fixation to vary while nodulation remained constant (blue) or allowing the rate of nodulation to vary while the rate of N_2_-fixation per gram nodule remained constant (red). The dashed lines indicate the maximal rate of plant growth with exogenous ammonium (upper) and the maximal rate of plant growth when relying on N_2_-fixation (lower).

We then simulated the effects of individual bacteria gene deletion on plant biomass production (Dataset S3) and compared the results to published experimental data. The model was able to accurately predict the phenotypes of many *S. meliloti* mutants. For example, *S. meliloti* genes such as *nifH* (nitrogenase), *dctA* (succinate transport), *ilvI* (branched chain amino acid biosynthesis), *aatA* (aspartate transaminase), *pgk* (phosphoglycerate kinase), and *nrdJ* (ribonucleotide reductase) were correctly predicted to be essential, while *pyc* (pyruvate carboxylase), *glnA* (glutamine synthetase), *pckA* (phosphoenolpyruvate carboxykinase), and *leuB* (leucine biosynthesis) were correctly predicted to be non-essential (74–81). Similarly, the removal of plant-encoded nodule sucrose synthase, phosphoenolpyruvate carboxylase, and homocitrate synthase reactions abolished nitrogen fixation, as expected (55, 82, 83). However, it is important to note that the predictions were not perfect. For example, deleting *argG* (arginine biosynthesis) or *carA* (carbamoyl phosphate synthase) did not result in the expected phenotypes, while the incorrect malic enzyme (*tme* instead of *dme*) was predicted to be essential (84–86). Taken together, these analyses provide support for the general reliability of ViNE as a representation of nodule metabolism.

### Metabolic progression and nutrient exchange during nodule development

The presence of five nodule zones in ViNE provided an opportunity to examine the metabolic changes associated with the development of an effective nodule. To accomplish this, FBA was used to predict the flux distribution through the integrated metabolic networks of each nodule zone, and to simulate the effects of individually deleting each gene, or removing each reaction, specifically in a single nodule zone. Additionally, a robustness analysis was performed to evaluate how perturbations in the flux of individual bacteroid reactions influence the predicted rate of plant growth. The outputs of these analyses are provided as Datasets S4 and S5 and summarized in Figures S1 and S2. For simplicity, here we focus on the reaction-level analyses, and we split the nodule into only three sections: uninfected (zone I), differentiating (zones IId, IIp, and IZ), and nitrogen-fixing (zone III) (Figure 4). Highlighting the overall similarity of zones IId, IIp, and IZ, and thus supporting their grouping, the robustness analysis indicated that roughly 90% of the bacteroid reactions that had to carry flux in one of these zones had to carry flux in all three zones to maximize plant growth.

**Figure 4.**
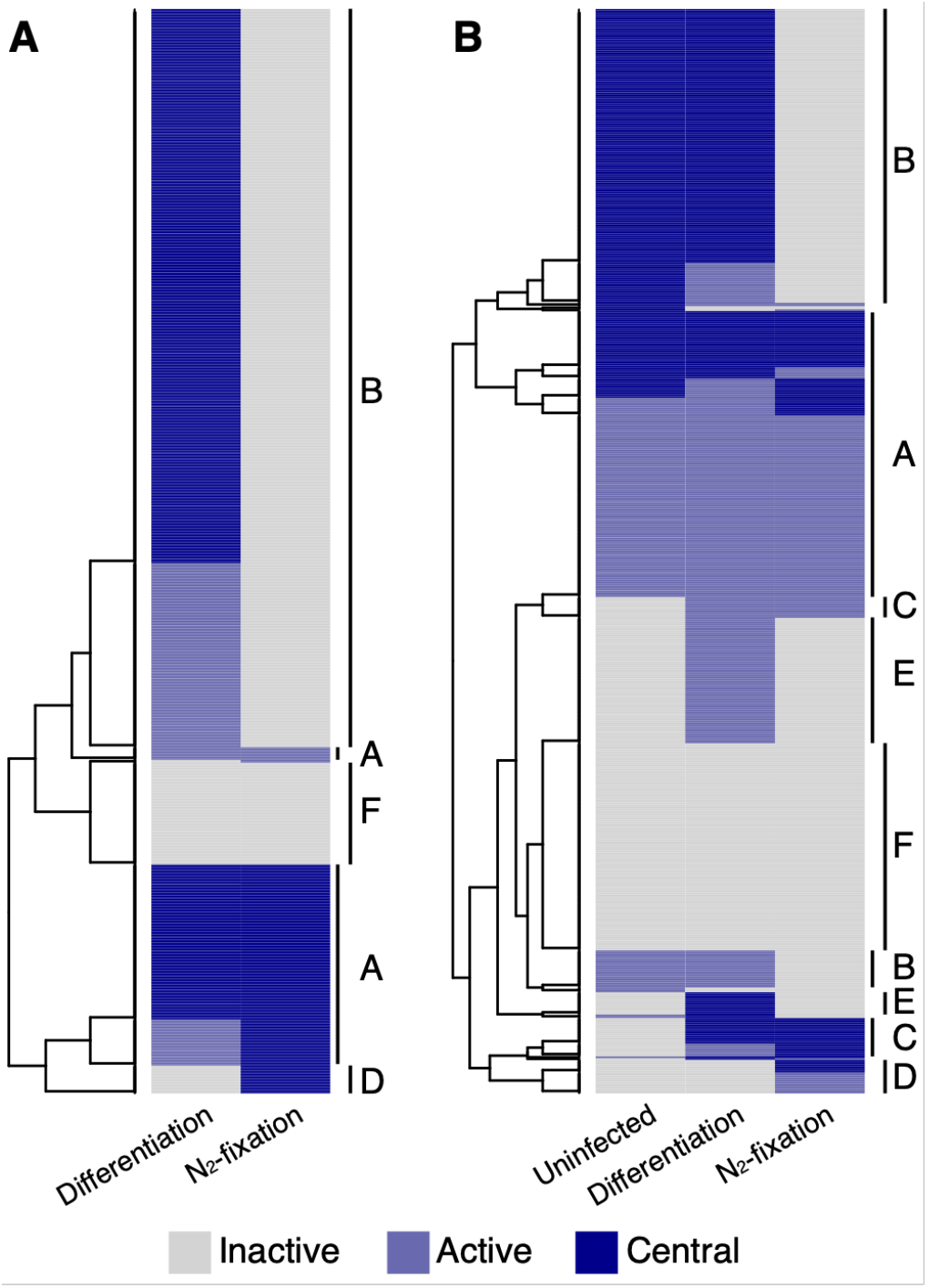
Nodule zone-specific metabolism. Heatmaps are presented displaying which reactions are inactive (grey), active but whose removal does not impair plant growth (light blue), or central (dark blue; growth reduction > 10% for the model missing the reaction compared to the full model) in the different nodule zones. In the differentiation zone, reactions are marked as central only if it was central in each of zone IId, IIp, and IZ. Heatmaps are shown for (**A**) *S. meliloti* reactions and (**B**) *M. truncatula* reactions. Reactions were clustered using hierarchical clustering, and the following main clusters were identified: A – constitutively active; B – specific to growing cells; C – specific to infected cells; D – specific to the nitrogen-fixation zone; E – specific to the differentiation zone; F – constitutively inactive.

The most notable difference comparing the uninfected and differentiation zones was an increase in the number of active reactions related to energy production, including carbon and nucleotide metabolism. This result suggests that the accommodation of differentiating bacteroids may place additional energy demands on the plant cell, and that few additional metabolic functions are required. In contrast, the transition from the differentiation zone to the nitrogen fixing zone was associated with a marked decrease in the number of active reactions in both the plant and bacterial cells, consistent with published transcriptomic and proteomic datasets (58, 87–89). Highlighting this result, ~ 560 bacteroid reactions had to carry flux in the differentiation zones to optimize plant growth, whereas only 167 bacteroid reactions had to carry flux in the N_2_-fixation zone for maximal plant growth.

The lack of biomass production in the N_2_-fixation zone meant that most biomass biosynthetic pathways were predicted to be inactive and non-essential. However, bacterial pathways related to the production of cofactors for nitrogenase or energy production remained essential; this included FMN, heme, cobalamin, pyridoxine phosphate, and glutathione biosynthesis, as well as the pentose phosphate pathway. Similarly, the TCA cycle, oxidative phosphorylation, and purine biosynthesis in bacteroids were predicted to be essential in the N_2_-fixation zone, presumably to supply the massive amounts of energy required by nitrogenase. Biosynthesis of methionine and SAM were also predicted to be essential. Few other notable bacterial reactions were required in the N_2_-fixation zone (Dataset S4). In the plant compartment, the majority of the active reactions were related to central carbon metabolism for the production of energy and C_4_-dicarboxylates for use by the bacteroids, while other active reactions were involved in the assimilation of ammonium through the formation of glutamine. Consistent with experimental works [reviewed by (18, 90)], the FBA results indicated that the plant nodule cells were provided sucrose as a carbon/energy source; in fact, ~ 30% of all carbon fixed by the plant leaves was sent to nodule zone III. The sucrose was then hydrolyzed and metabolized to phosphoenolpyruvate, of which ~ 80% was diverted to oxaloacetate through a cytoplasmic phosphoenolpyruvate carboxykinase reaction for use in the production of C_4_-dicarboxylates.

Next, nutrient exchange between the plant and bacterial partners was examined. While the prevailing evidence suggests C_4_-dicarboxlyates (succinate, malate, fumarate) are the primary carbon source for N_2_-fixing bacteroids (18, 91–93), the source of carbon for differentiating bacteroids has not been established. The FBA results suggested that differentiating bacteroids primarily use sugars, likely sucrose, as a carbon source. This is consistent with micrographic evidence suggesting that bacterial mutants unable to use C_4_-dicarboxylates can undergo at least partial differentiation (91, 92). Currently, it is commonly accepted that nitrogen is primarily exported from bacteroids as ammonia (94, 95); however, some studies have suggested that L-alanine could be a major nitrogen export product (96, 97). Here, the FBA simulations were consistent with ammonia being the primary export product in the *S. meliloti* – *M. truncatula* symbiosis. However, prior to constraining the nodule reaction space, reducing the availability of oxygen to the bacteroids resulted in a shift in the nitrogen export product from ammonia to L-alanine. Thus, the detection of L-alanine versus ammonia as an export product could be due, in part, to differences in experimental set-up that may influence oxygen availability to the bacteroid. Also, experimental data suggest that rhizobial biosynthesis of some amino acids are essential for the symbiosis while the biosynthesis of others are not, and that the phenotypes may be symbiosis-specific [reviewed by (98)]. Similarly, our FBA simulations suggested that rhizobial biosynthesis of approximately half of the amino acids was essential for the symbiosis.

### Sociobiology of symbiotic nitrogen fixation

By containing a representation of an entire nodule, ViNE allowed for an evaluation of the metabolic costs associated with SNF. Using FBA, the maximal plant growth rate of the nodulated system (without exogenous ammonium) was estimated to be ~ 72% of the maximal growth rate of a nodule-free system supplied with non-limiting amounts of exogenous ammonium (Figure 3, Table 2). The largest factor contributing to the difference in growth was the direct energetic cost of supporting N_2_-fixation (~ 67% of the difference; Table 2). The remaining third of the difference was explained by the cost of synthesizing (~ 11% of the difference) and maintaining (~ 22% of the difference) the nodule and bacteroid tissue (Table 2).

**Table 2.**
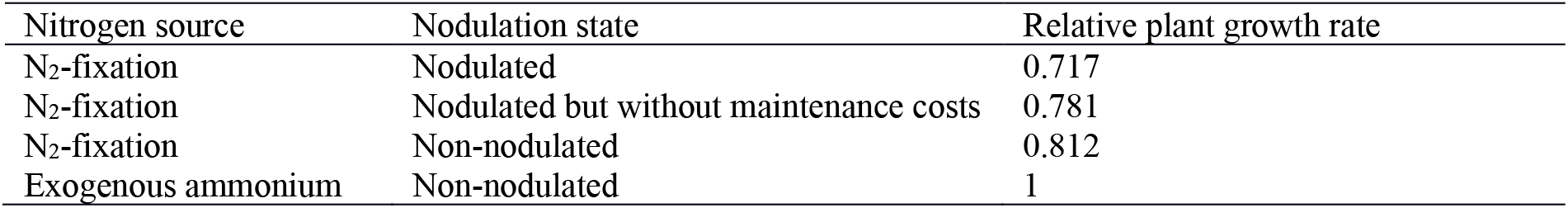
Contributions of N_2_-fixation and nodulation to the fitness costs of SNF.

We next evaluated the relationship between the rate of N_2_-fixation (without modifying the plant to nodule ratio) and the rate of plant growth. When the rate of N_2_-fixation was below the optimum, there was a linear relationship between N_2_-fixation and biomass production (Figure 5A). However, excessive N_2_-fixation quickly resulted in impaired plant growth, with a 10% excess of N_2_-fixation collapsing the symbiosis (Figure 5A). We hypothesized that this result was due to insufficient energy to support both the excess N_2_-fixation and the ATP maintenance costs. Consistent with this hypothesis, removing the upper limit on the rate of zone III oxygen uptake resulted in a gradual decrease in plant growth as the rate of N_2_-fixation was increased above the optimal (Figure 5A). In this case, excessive N_2_-fixation was less detrimental than insufficient N_2_-fixation; the effect of increasing the rate of N_2_-fixation by 1 µmol hr^−1^ (g plant dry weight)^−1^ increased or decreased the rate of plant growth by 14.7 or 3.4 mg day^−1^ (g plant dry weight)^−1^ when below or above the optimum, respectively. We next examined the consequences of varying the rate of nodulation (i.e., the ratio between plant and nodule biomass) while maintaining a constant rate of N_2_-fixation per gram of nodule. The simulations demonstrated linear relationships between the rate of nodulation and plant growth both above and below the optimum (Figure 5B), with increasing the percent nodulation resulting in a 3-fold greater impact when below the optimum compared to above the optimum. Overall, these simulations suggest that a slightly too efficient symbiosis is preferable (for plant biomass production) over a slightly inefficient symbiosis, unless the required rate of O_2_ usage exceeds the nodule oxygen diffusion limit.

**Figure 5.**
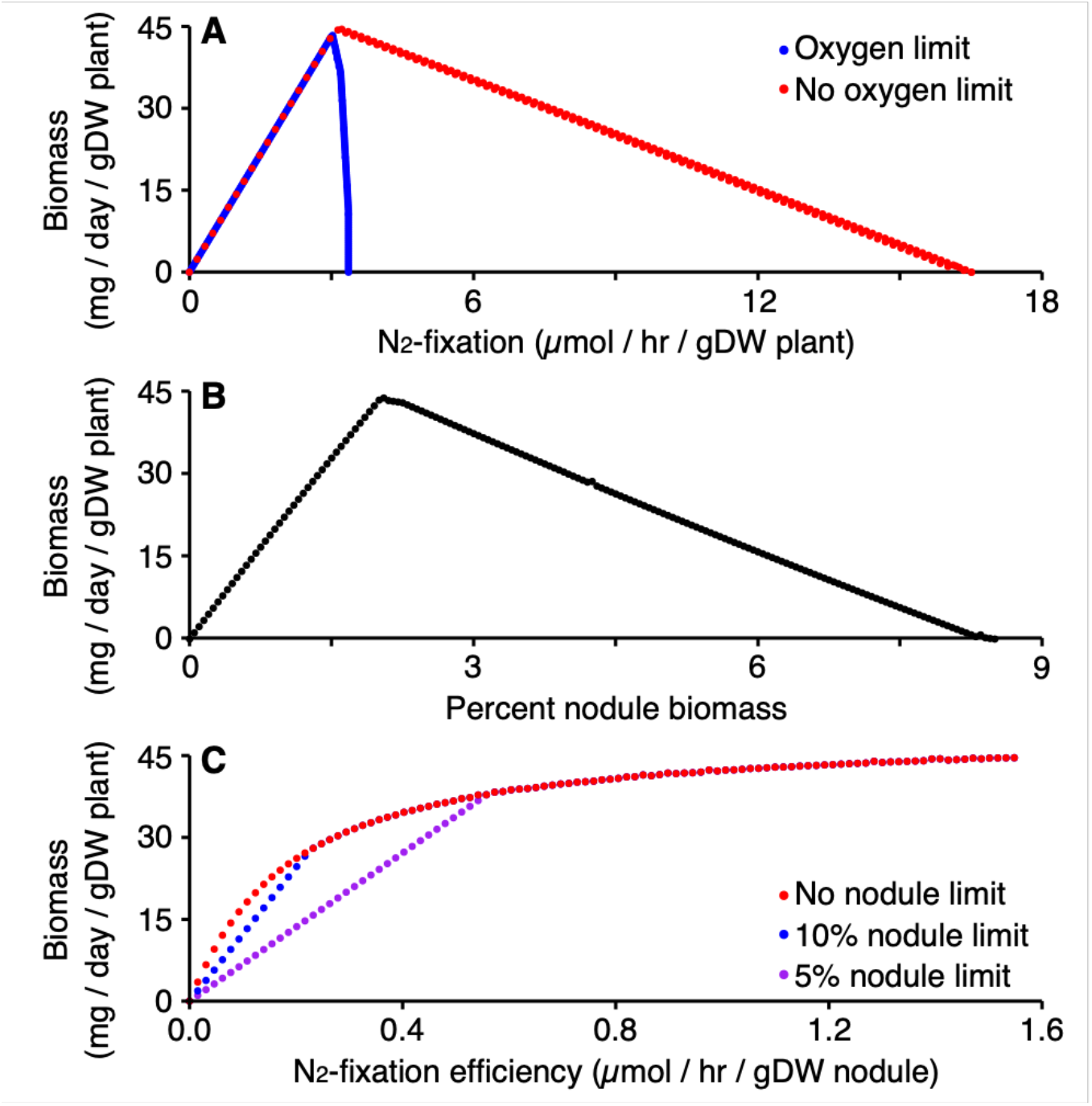
Relations between plant growth and rate of N_2_-fixation or nodulation. (**A**) Pareto frontiers showing the relationship between the rate of N_2_-fixation (with a constant ratio between plant and nodule biomass) and the rate of plant biomass production using ViNE with default parameters (blue) or no limit on zone III oxygen usage (red). (**B**) The relationship between the amount of nodule per plant, expressed as a percent of total (plant + nodule) biomass (with a constant rate of N_2_-fixation per gram of nodule) and the rate of plant biomass production. (**C**) The effect of N_2_-fixation efficiency (rate of N_2_-fixation per gram nodule) on the rate of plant growth, with the amount of nodule biomass optimized to maximize plant growth and without a limit on zone III oxygen uptake (see Figure S3 for simulations with an oxygen uptake limit). Nodule biomass was either uncapped (red) or limited to 10% (blue) or 5% (purple) of the overall biomass.

The previous simulations represented simple scenarios where only a single variable differed. In reality, a change in the efficiency of N_2_-fixation should be accompanied by a change in the extent of nodulation as a result of legume autoregulation of nodulation (99). We therefore ran simulations where the efficiency of N_2_-fixation (i.e., the rate of N_2_-fixation per gram of nodule) was varied and the amount of nodule biomass was optimized to maximize plant growth. Strikingly, the simulations suggested a pattern of diminishing returns associated with increasing the efficiency of N_2_-fixation (Figures 5C and S3); decreasing N_2_-fixation efficiency 50% from the maximum tested value resulted in a mere 10% decrease in plant growth. The half maximal growth rate was achieved with a N_2_-fixation efficiency of just 10% the maximal, although this required that the nodule accounted for almost 13% of the total biomass. If we assume an upper limit of nodulation at 10% or 5% of the total biomass, the benefits of low rates of N_2_-fixation are decreased although the pattern of diminishing returns remains (Figures 5C and S3). In these cases, half maximal plant growth rate is achieved at 12% or 21%, respectively, of the highest tested N_2_-fixation efficiency. Overall, these simulations support that even a poor symbiosis is likely to provide a noticeable benefit to the plant.

### Influence of protons and O_2_ on the carbon source provided to N_2_-fixing bacteroids

It is well-established that C_4_-dicarboxylates (malate, succinate, fumarate) are the primary carbon source provided to nitrogen-fixing zone III bacteroids (18, 91–93); however, the reason for this remains unclear. We therefore attempted to uncover a metabolic explanation using ViNE. Surprisingly, preliminary FBA simulations with the ViNE precursor model (i.e., prior to constraining the reaction space) suggested that the N_2_-fixing bacteroids of zone III use sucrose, not C_4_-dicarboxylates, as the primary carbon source. Unexpectedly, forcing the use of C_4_-dicarboxylates resulted in the model being unable to fix nitrogen or produce plant biomass. During those simulations, protons of the plant cytosol could be transferred to the peribacteroid space but were not allowed to be used by the N_2_-fixing bacteroids. However, this may not be realistic since the peribacteroid space of N_2_-fixing bacteroids is acidic due to import of protons from the plant cytosol (100–102). If the analysis was repeated and the zone III bacteroids were provided access to the protons of the peribacteroid space, it became possible for C_4_-dicarboxylates to serve as the primary carbon source and support N_2_-fixation and plant growth. These results suggest that the plant-driven acidification of the peribacteroid space is essential for the metabolic functioning of the bacteroid.

Although the transfer of protons to the periplasm allowed C_4_-dicarboxylates to support N_2_-fixation, the rate of plant biomass production nevertheless remained higher when the N_2_-fixing bacteroids were provided sucrose instead of C_4_-dicarboxylates. To further investigate this difference, ViNE was modified to contain reactions for the transport and metabolism of sucrose by N_2_-fixing bacteroids (see Materials and Methods). Consistent with results from the precursor model, FBA simulations suggested the ability of bacteroids to use sucrose (plus C_4_-dicarboxylates) increased plant growth rate by 6.4% relative to when bacteroids were supplied only C_4_-dicarboxylates. ViNE contains a limit on the rate of oxygen uptake by zone III nodule tissue, restricting nodule and bacteroid metabolism. We wondered whether the use of sucrose versus C_4_-dicarboxylates may be modulated by the free oxygen concentration of the nodule. The concentration of free oxygen in the N_2_-fixation zone has been experimentally demonstrated to be < 50 nM (103). Notably, the K_m_ values of the mitochondrial and bacterial terminal oxidases towards oxygen are 50-100 nM (104, 105) and 7 nM (106), respectively. These enzyme kinetics suggest that the metabolism of the plant fraction, but not the bacteroid fraction, of the nodule is likely to be oxygen limited (107, 108), a conclusion that is supported by measurements of nodule adenylate pools (109). Therefore, we ran a series of simulations in which the upper limit of the mitochondrial terminal oxidase reaction of zone III was varied, with no overall limit on the use of oxygen by the nodule. Gradually reducing the flux through the mitochondrial terminal oxidase was associated with a gradual replacement of sucrose with C_4_-dicarboxylates as the bacteroid carbon source (Figure 6). This result is consistent with the hypothesis that the low free oxygen concentration of the N_2_-fixation zone could be a contributing factor to why bacteroids are provided C_4_-dicarboxylates, and not sugars, as the primary carbon source.

**Figure 6.**
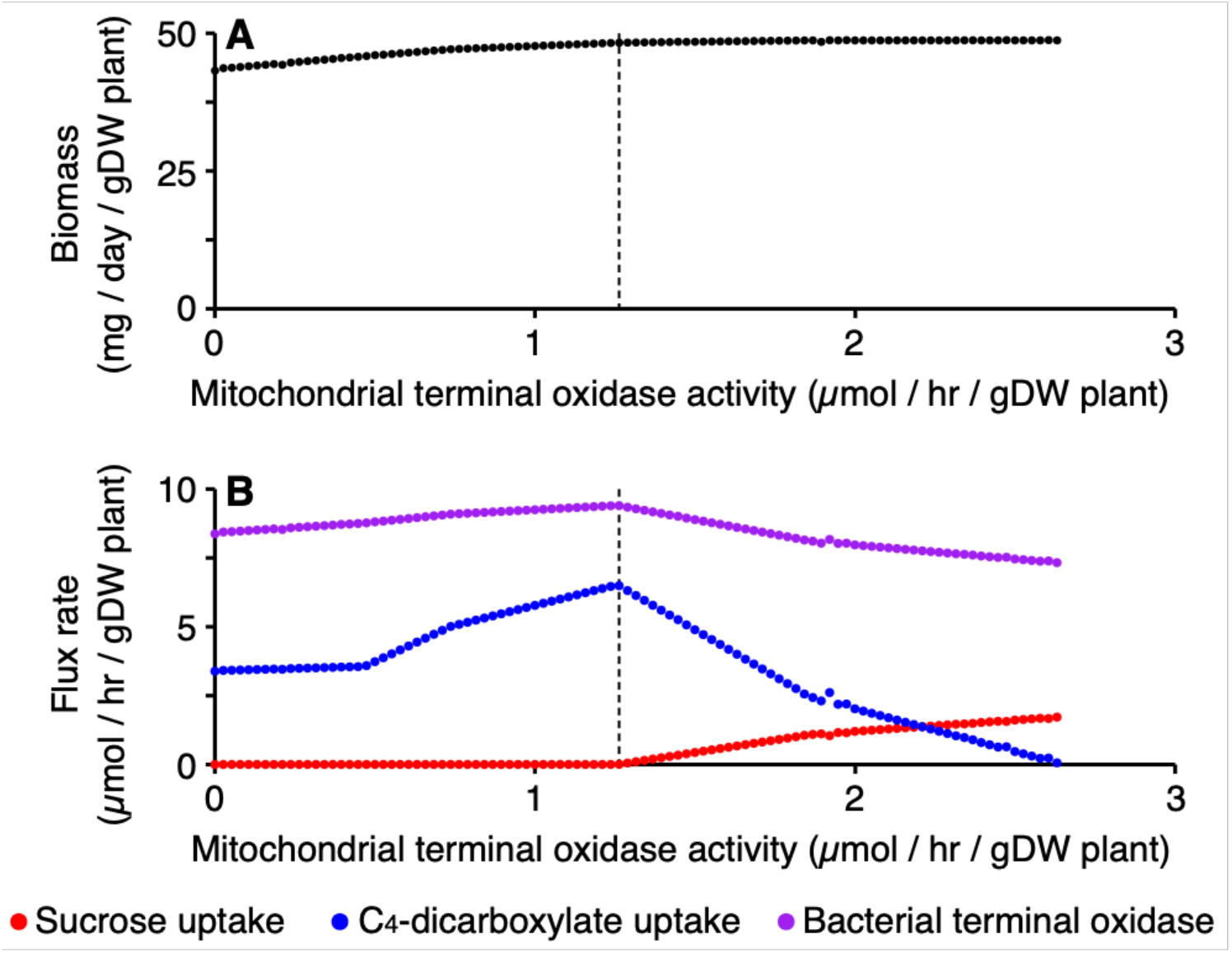
Effects of limiting the mitochondrial terminal oxidase in the N_2_-fixing zone. FBA simulations were run with a modified version of ViNE in which bacteroid metabolism can be supported by sucrose in addition to C_4_-dicarboxylates. No overall limit on oxygen usage of the nodule was set during the simulations, but a limit was set on the activity of the mitochondrial terminal oxidase of the zone III nodule tissue. (**A**) The effect on plant growth rate of varying the mitochondrial terminal oxidase of the zone III nodule tissue. (**B**) The effect on specified flux rates of varying the mitochondrial terminal oxidase of the zone III nodule tissue. Red – the flux rate of sucrose uptake by N_2_-fixing bacteroids; blue – the flux rate of C_4_-dicarboxylates of N_2_-fixing bacteroids; purple – the flux rate of the terminal oxidase of N_2_-fixing bacteroids. The dashed line indicates the mitochondrial terminal oxidase flux rate below which no sucrose is used by the bacteroids.

Assuming the nodule (consisting primarily of zone III tissue) accounts for 2% of total plant biomass, and that bacteroid biomass accounts for 25% of nodule biomass, the maximal rate of predicted C_4_-dicarboxylate import by N_2_-fixing bacteroids (1.3 mmol hr^−1^ [g bacteroid dry weight]^−1^) was similar to experimentally determined uptake rates by *S. meliloti* bacteroids (1.1 to 1.3 mmol hr^−1^ [g bacteroid dry weight]^−1^) (35, 110). Considered together, these simulations provide evidence that C_4_-dicarboxylates can support optimal plant growth under physiologically-relevant conditions.

## DISCUSSION

Models of the integrated metabolism of various holobionts (consisting of a host and its symbiotic microorganisms) would be valuable tools to understand the emergent properties of these systems (111, 112). However, to date there are few examples of constraint-based metabolic modelling being used to study metabolic interactions [e.g., (8, 31)], with this approach most commonly used to study the human gut microbiome (113). Here, we developed a broadly adaptable pipeline for modelling the metabolism of interacting organisms across physiologically distinct tissue (sub)sections. Using metabolic network reconstruction and constraint-based modelling, we studied the metabolism of a legume root nodule and SNF, a well-established model of inter-organismal metabolic exchange and cellular differentiation. Our model (ViNE) accounts for plant shoot, root, and nodule tissues, with the nodule encompassing the metabolism of both the plant and bacterial partners and subdivided into five developmental zones. This is an advance over previous attempts at modelling SNF (33, 35–40), most of which focused solely on bacterial metabolism while treating the plant as a black box. The increased complexity of ViNE allows for more accurate simulations of the nutrient exchange, analysis of the metabolic differentiation associated with nodule development, examination of unexpected emergent properties of the symbiosis resulting from inter-organism interactions, and for the possibility to perturb the network at the single reactions level. Initial simulations with ViNE supported that this model does a good job at capturing the metabolism of a legume nodule. Nevertheless, as with all models, ViNE predictions were imperfect. However, as we often compared simulated phenotypes for *M. truncatula* with experimental data for *M. sativa*, and given that rhizobium mutant phenotypes are often plant specific [e.g., (114–116)], we cannot rule out that some of the inconsistencies are the result of plant-specific phenotypes. Going forward, we intend to continue to manually refine and update ViNE to maximize consistency with experimental observations.

FBA simulations with ViNE revealed a pattern of diminishing returns in terms of plant growth (as a proxy for fitness) as the efficiency of the symbiosis (rate of N_2_-fixation per gram of nodule) increased, assuming that the amount of nodule biomass per plant could vary (Figure 5C). This observation has potential implications for engineering SNF for biotechnological applications. It suggests that when developing rhizobial inoculants, maximizing competition for nodule occupancy may have a greater impact than maximizing the rate of N_2_-fixation. This result also supports efforts aimed at engineering N_2_-fixing symbiosis with cereals (117) by highlighting how even a low efficiency symbiosis has the potential to have a noticeable benefit on crop yield.

At the same time, the pattern of diminishing returns is interesting from an evolutionary perspective (118). In particular, the evolution of N_2_-fixation efficiency may be influenced by the rhizobium community diversity, assuming that nodule infection increases the fitness of rhizobia (119). In an environment dominated by a single rhizobium, kin selection may favour the evolution of a poorly efficient symbiosis as it would increase nodule number and thus the size of the niche for colonization by the rhizobia. On the other hand, in a highly diverse environment, evolution of strains capable of entering into a highly efficient symbiosis may be favoured, as this would lead to fewer nodules and thus less plant resources being allocated to competing rhizobium strains, thereby limiting the spread of less mutualist (viz. cheater) strains (20, 120).

Of particular interest to us was the metabolic exchange between the plant and rhizobia, both during N_2_-fixation and during differentiation. The carbon source(s) of differentiating rhizobia remain poorly understood. Results with ViNE suggested that sucrose may be a major carbon source for the differentiating bacteroids. However, *S. meliloti* mutants unable to transport sucrose are not impaired in nodule formation (121), suggesting that differentiating bacteroids have access to at least one other carbon source. Interestingly, a *S. meliloti pyc* mutant unable to grow with glycolytic carbon sources was not impaired in differentiation (80). Similarly, *S. meliloti pckA* (78) and *tpi* (122) mutants unable to grow with gluconeogenic carbon sources remained capable of differentiating. Thus, it seems likely that differentiating bacteroids have access to a variety of glycolytic and gluconeogenic carbon substrate, with sugars possibly serving as the main carbon source in wild type nodules. If so, the restriction of carbon flow to N_2_-fixing bacteroids to just C_4_-dicarboxylates may be the result of active remodelling of the peribacteroid membrane during differentiation.

In attempting to identify conditions favouring the use of C_4_-dicarboxylates as a carbon source by N_2_-fixing bacteroids, ViNE also provided insights into the metabolic exchange in the N_2_-fixation zone. The peribacteroid space of N_2_-fixing bacteroids is known to be acidic due to the activity of H^+^-ATPases on the peribacteroid membrane (100–102). This acidification contributes to the import of C_4_-dicarboxylates and the export of ammonium from/to the plant cytosol and the peribacteroid space (123), and it may contribute to the lysis of non-functional symbiosomes (124). Our FBA simulations suggest that the plant-derived protons of the peribacteroid space may also be actively used by the bacteroid to support its metabolism.

Although it is generally accepted that nodules are low oxygen environments (103), the site of O_2_-limitation has been debated. Based on the average concentration of free oxygen in the nodule, enzyme kinetics data are consistent with the mitochondria being O_2_-limited and the bacteroids being O_2_-sufficient (103–106). Measurements of the adenylate pools of the plant and bacterial nodule fractions support this conclusion (109). However, others have argued that nodule adenylate measurements suggest that bacteroids, not the plant, are O_2_-limited (125). Similarly, it was suggested that mitochondria cluster near the periphery of the cell near air pockets, resulting in elevated local O_2_ concentrations (126, 127). The FBA results presented here predicted that C_4_-dicarboxylates are the optimal carbon source for N_2_-fixing bacteroids only when the plant mitochondria are O_2_-limited while the bacteroids are O_2_-sufficient (Figure 6). This result supports the hypothesis that mitochondria, and not bacteroids, are O_2_-limited in wild type nodules.

In sum, this work presented a complex metabolic model representing the full metabolism of a rhizobium-nodulated legume, as well as a series of simulations demonstrating the potential for this model to help address genetic, evolutionary, metabolic, and sociobiological questions. Future work will be aimed at continuing to refine and improve the quality of the model, and to using the model to generate hypotheses to guide experimental studies and to assist in the interpretation of experimental datasets.

## ACKNOWLEDGEMENTS

GCD was supported by a postdoctoral fellowship from the Natural Science and Engineering Research Council of Canada, and funding from Queen’s University. AM was supported by a grant from Fondazione Cassa di Risparmio di Firenze (project name: Metatrack)

## SUPPLEMENTARY MATERIAL

**Supplementary File S1:** Additional text, figures and tables.

**Text S1.** Simulations involving varying rates of nodulation.

**Table S1.** Biomass composition of iGD1348.

**Table S2.** Summary properties of the *S. meliloti* metabolic reconstruction iGD1348.

**Figure S1.** Nodule zone-specific analysis of essential metabolism.

**Figure S2.** Bacteroid robustness analysis summary.

**Supplementary File S2:** SBML, XLS, and MATLAB COBRA formats of the *S. meliloti* updated metabolic reconstruction (iGD1348) used to generate the ViNE.

**Supplementary File S3:** SBML, XLS, and MATLAB COBRA formats of the ViNE metabolic reconstruction.

**Dataset S1:** The list of the reactions excluded from the model following dead-end metabolites removal

**Dataset S2:** Comparison of experimental and predicted *S. meliloti* growth phenotypes.

**Dataset S3:** The effects of individual bacteria gene deletion on plant biomass production

**Dataset S4:** Nodule zone-specific gene deletion and reaction removal analyses

**Dataset S5:** Bacteroid robustness analysis.

