## Supplementary Material S1 for "A Virtual Nodule Environment (ViNE) for modelling the inter-kingdom metabolic integration during symbiotic nitrogen fixation"

**Text S1. Simulations involving varying rates of nodulation.**

The following strategy was used for simulations were $NH_4$ was added to the soil and the rate of nodulation (i.e., gram of nodule per gram of plant) was optimized to maximize plant growth while the efficiency of $N_2$-fixation (i.e., mmol of $N_2$ fixed per gram of nodule) remained constant. First, the model was optimized in the absence of exogenous $NH_4$. Next, nodule biomass was removed from the overall biomass reaction and instead forced through a sink reaction at a rate equal to its synthesis in the previous step. The desired rate of $NH_4$ availability was then set and plant biomass production optimized. Based on the rates of plant growth and $N_2$-fixation in the previous step, and the desired efficiency of nitrogen-fixation, the rate that nodule biomass was produced was updated. Plant biomass production was again optimized, and the ratio between plant and nodule biomass production was used to update all nodule maintenance cost reactions. The following loop was then employed. Plant biomass production was optimized and the rates of plant growth and nitrogen-fixation, together with the desired efficiency of nitrogen-fixation, were used to update the rate of nodule biomass production. Plant biomass production was again optimized and the ratio between plant and nodule biomass production was used to update all nodule maintenance cost reactions. Once again, plant biomass production was optimized, and the maximal rate of zone III oxygen uptake was set based on the ratio between nodule and plant biomass production and the desired oxygen limit. Plant biomass production was optimized, and the $N_2$-fixation efficiency was calculated. This loop was iterated until the $N_2$-fixation efficiency was within 0.1% of the desired value, at which point the results were recorded.

The following strategy was used for simulations where the efficiency of $N_2$-fixation (i.e., mmol of $N_2$ fixed per gram of nodule) was varied while the rate of nodulation (i.e., gram of nodule per gram of plant) was optimized to maximize plant growth. First, the model was optimized with default settings, following which nodule biomass was removed from the overall biomass reaction and instead forced through a sink reaction at a rate equal to its synthesis in the initial FBA solution. The following loop was then employed. The desired efficiency of $N_2$-fixation was set, and the rate of nodule biomass production updated based on the difference in efficiency relative to default settings (i.e., if efficiency was reduced by 50%, nodule biomass was doubled). Additionally, the maximal rate of zone III oxygen uptake and $N_2$-fixation was adjusted based on the updated ratio between nodule and plant biomass production. The rate of plant biomass production was then optimized. Based on the rate of $N_2$-fixation in the solution of the previous step, the rate of biomass production was updated, the lower limits of all nodule maintenance cost reactions were updated, and the maximal rate of zone III oxygen uptake and $N_2$-fixation were updated. The rate of plant biomass production was again optimized. This loop was iterated until the $N_2$-fixation efficiency was within 0.5% of the desired value. We found that this loop only identified local maximums for the rate of plant biomass production. In order to find the global maximum, a second loop was employed. In each loop, the rate of nodulation was slightly increased relative to the previous solution. Next, the previous loop was again employed to identify the local maximum. This was repeated until the local maximum following a slight increase in the rate of nodulation was lower than the previous local maximum; we assume this meant the previous iteration represented the global maximum. The results for the global maximum were then recorded.

To examine the effect of $N_2$-fixation efficiency on plant growth in the absence of a zone III oxygen limitation, the same process was employed with the exception that the maximal rate of zone III oxygen uptake was kept constant at 1,000 µmol $hr^{-1}$ (g plant dry weight)$^{-1}$. In cases where the rate of nodulation was capped at 5% of 10%, the same process was used with the exception that the rate of nodule biomass production was limited to 5% or 10% the sum of plant and nodule biomass production.

**Table S1.** Biomass composition of iGD1348.

| Component | % dry mass | Composition - % | | | |
|---|---|---|---|---|---|
| DNA [a] | 3.56 | Guanine - 31.05 | Adenine - 18.95 | Cytosine - 31.05 | Thymine - 18.95 |
| RNA [b] | 9.03 | Guanine - 28.09 | Adenine - 21.91 | Cytosine - 28.09 | Uracil - 21.91 |
| Protein [c] | 62.72 | Lysine - 3.20 | Alanine - 12.01 | Phenylalanine - 3.94 | Leucine 10.19 |
| | | Arginine 7.33 | Glutamine - 2.90 | Glycine - 8.46 | Methionine - 2.44 |
| | | Valine - 7.58 | Proline - 5.03 | Tyrosine - 2.29 | Aspartate - 5.31 |
| | | Glutamate - 5.84 | Histidine - 2.11 | Threonine - 5.15 | Isoleucine - 5.48 |
| | | Cysteine - 0.93 | Asparagine - 2.64 | Tryptophan - 1.38 | Serine - 5.80 |
| Phosphatidylglycerol [d] | 1.274 | Phosphatidylglycerol-1-palmitoleoyl-2-palmitic - 0.4 | | | |
| | | Phosphatidylglycerol-1,2-palmitic-2-palmitic - 1.4 | | | |
| | | Phosphatidylglycerol-1-cis-vaccenoyl-2-palmitoleoyl - 1.8 | | | |
| | | Phosphatidylglycerol-1-cis-vaccenoyl-2-palmitic - 24.9 | | | |
| | | Phosphatidylglycerol-1-palmitic-2-cis-vaccenoyl - 1.7 | | | |
| | | Phosphatidylglycerol-1-cis-vaccenoyl-2-palmitoleoyl(cyclopropanated) - 1.1 | | | |
| | | Phosphatidylglycerol-1-cis-vaccenoyl(cyclopropanated)-2-palmitoleoyl - 1.1 | | | |
| | | Phosphatidylglycerol-1-cis-vaccenoyl(cyclopropanated)-2-palmitic - 2.6 | | | |
| | | Phosphatidylglycerol-1-cis-vaccenoyl-2-cis-vaccenoyl - 57.3 | | | |
| | | Phosphatidylglycerol-1-cis-vaccenoyl(cyclopropanated)-2-cis-vaccenoyl - 7.3 | | | |
| | | Phosphatidylglycerol-1-cis-vaccenoyl(cyclopropanated)-2-cis-vaccenoyl(cyclopropanated) - 0.3 | | | |
| Cardiolipin [d] | 0.506 | Cardiolipin-1,2-(1-palmitoleoyl-2-palmitic) - 0.4 | | | |
| | | Cardiolipin-1,2-(1,2-palmitic) - 1.4) | | | |
| | | Cardiolipin-1,2-(1-cis-vaccenoyl-2-palmitoleoyl) - 1.8 | | | |
| | | Cardiolipin-1,2-(1-cis-vaccenoyl-2-palmitic) - 24.9 | | | |
| | | Cardiolipin-1,2-(1-palmitoleoyl-2-palmitic) - 1.7 | | | |
| | | Cardiolipin-1,2-(1-cis-vaccenoyl-2-palmitoleoyl(cyclopropanated)) - 1.1 | | | |
| | | Cardiolipin-1,2-(1-cis-vaccenoyl(cyclopropanated)-2-palmitoleoyl) - 1.1 | | | |
| | | Cardiolipin-1,2-(1-cis-vaccenoyl(cyclopropanated)-2-palmitic) - 2.6 | | | |
| | | Cardiolipin-1,2-(1-cis-vaccenoyl-2-cis-vaccenoyl) - 57.3 | | | |
| | | Cardiolipin-1,2-(1-cis-vaccenoyl(cyclopropanated)-2-cis-vaccenoyl) - 7.3 | | | |
| | | Cardiolipin-1,2-(1-cis-vaccenoyl(cyclopropanated)-2-cis-vaccenoyl(cyclopropanated)) - 0.3 | | | |
| Phosphatidylethanolamine [d] | 2.232 | Phosphatidylethanolamine-1-cis-vaccenoyl-2-palmitoleoyl - 0.9 | | | |
| | | Phosphatidylethanolamine-1-cis-vaccenoyl-2-palmitic - 7.0 | | | |

| | | |
|---|---|---|
| | | Phosphatidylethanolamine-1-cis-vaccenoyl-2-palmitoloeyl(cyclopropanated) - 1.05 |
| | | Phosphatidylethanolamine-1-cis-vaccenoyl(cyclopropanated)-2-palmitoloeyl - 1.05 |
| | | Phosphatidylethanolamine-1-cis-vaccenoyl(cyclopropanated)-2-palmitic - 2.9 |
| | | Phosphatidylethanolamine-1,2-cis-vaccenoyl - 47.2 |
| | | Phosphatidylethanolamine-1-cis-vaccenoyl(cyclopropanated)-2-cis-vaccenoyl - 35.5 |
| | | Phosphatidylethanolamine-1-cis-vaccenoyl(cyclopropanated)-2-cis-vaccenoyl(cyclopropanated) - 4.4 |
| Monomethylethanolamine [d] | 1.897 | Monomethyl-phosphatidylethanolamine-1-cis-vaccenoyl-2-palmitoleoyl - 0.9 |
| | | Monomethyl-phosphatidylethanolamine-1-cis-vaccenoyl-2-palmitic - 7.0 |
| | | Monomethyl-phosphatidylethanolamine-1-cis-vaccenoyl-2-palmitoleoyl(cyclopropanated) - 1.05 |
| | | Monomethyl-phosphatidylethanolamine-1-cis-vaccenoyl(cyclopropanated)-2-palmitoleoyl - 1.05 |
| | | Monomethyl-phosphatidylethanolamine-1-cis-vaccenoyl(cyclopropanated)-2-palmitic - 2.9 |
| | | Monomethyl-phosphatidylethanolamine-1,2-cis-vaccenoyl - 47.2 |
| | | Monomethyl-phosphatidylethanolamine-1-cis-vaccenoyl(cyclopropanated)-2-cis-vaccenoyl - 35.5 |
| | | Monomethyl-phosphatidylethanolamine-1-cis-vaccenoyl(cyclopropanated)-2-cis-vaccenoyl(cyclopropanated) - 4.4 |
| Phosphatidylcholine [d] | 9.758 | Phosphatidylcholine-1-cis-vaccenoyl-2-palmitoleoyl - 0.9 |
| | | Phosphatidylcholine-1-cis-vaccenoyl-2-palmitic - 7.0 |
| | | Phosphatidylcholine-1-cis-vaccenoyl-2-palmitoleoyl(cyclopropanated) - 1.05 |
| | | Phosphatidylcholine-1-cis-vaccenoyl(cyclopropanated)-2-palmitoleoyl - 1.05 |
| | | Phosphatidylcholine-1-cis-vaccenoyl(cyclopropanated)-2-palmitic - 2.9 |
| | | Phosphatidylcholine-1,2-cis-vaccenoyl - 47.2 |
| | | Phosphatidylcholine-1-cis-vaccenoyl(cyclopropanated)-2-cis-vaccenoyl - 35.5 |
| | | Phosphatidylcholine-1-cis-vaccenoyl(cyclopropanated)-2-cis-vaccenoyl(cyclopropanated) - 4.4 |
| Sulfoquinovosyldiacylglycerol [d] | 0.326 | Sulfoquinovosyl-1-palmitoleoyl-2-palmitic-sn-glycerol - 3.0 |
| | | Sulfoquinovosyl-1,2-palmitic-sn-glycerol - 17.2 |
| | | Sulfoquinovosyl-1-cis-vaccenoyl-2-palmitoleoyl - 1.4 |
| | | Sulfoquinovosyl-1-cis-vaccenoyl-2-palmitic - 21.8 |
| | | Sulfoquinovosyl-1-palmitic-2-cis-vaccenoyl - 10.1 |
| | | Sulfoquinovosyl-1-cis-vaccenoyl-2-palmitoleoyl(cyclopropanated) - 0.75 |
| | | Sulfoquinovosyl-1-cis-vaccenoyl(cyclopropanated)-2-palmitoleoyl - 0.75 |
| | | Sulfoquinovosyl-1-cis-vaccenoyl(cyclopropanated)-2-palmitic - 3.1 |
| | | Sulfoquinovosyl-1-palmitic-2-cis-vaccenoyl(cyclopropanated)-sn-glycerol - 2.3 |
| | | Sulfoquinovosyl-1-cis-vaccenoyl-2-cis-vaccenoyl-sn-glycerol - 17.8 |
| | | Sulfoquinovosyl-1-cis-vaccenoyl(cyclopropanated)-2-cis-vaccenoyl-sn-glycerol - 10.5 |
| | | Sulfoquinovosyl-1-cis-vaccenoyl(cyclopropanated)-2-cis-vaccenoyl(cyclopropanated)-sn-glycerol - 0.5 |
| | | Sulfoquinovosyl-1-cis-vaccenoyl-steric-sn-glycerol - 10.8 |

| | | |
|---|---|---|
| Ornithine lipids [d] | 0.293 | Ornithine-1-palmitic-2-cis-vaccenoyl - 0.5<br>Ornithine-1-steric-2-palmitoleoyl - 0.7<br>Ornithine-1-palmitic-2-cis-vaccenoyl(cyclopropanated) - 1.0<br>Ornithine-1-steric-2-palmitoleoyl(cyclopropanated) - 2.1<br>Ornithine-1-cis-vaccenoyl-2-cis-vaccenoyl - 3.0<br>Ornithine-1-steric-2-cis-vaccenoyl - 37.1<br>Ornithine-1-cis-vaccenoyl-2-cis-vaccenoyl(cyclopropanated) - 5.3<br>Ornithine-1-steric-2-cis-vaccenoyl(cyclopropanated) - 49.8<br>Ornithine-1-cis-vaccenoyl(cyclopropanated)-2-cis-vaccenoyl(cyclopropanated) - 0.6 |
| Poly-3-hydroxybutyrate | 1 | N/A |
| Glycogen | 0.1 | N/A |
| Lipopolysaccharide | 3.82 | N/A |
| Peptidoglycan | 2.54 | N/A |
| Low molecular weight succinoglycan [e] | 0.4 | N/A |
| High molecular weight succinoglycan [e] | 0.1 | N/A |
| Putrescine | Trace | N/A |
| Spermidine | Trace | N/A |
| Vitamins, cofactors, coenzymes, ions, and other [g] | Trace | Polyphosphate<br>Pantothenate<br>Coenzyme A<br>$NAD^+$; NADH<br>$NADP^+$; NADPH<br>$FAD^+$, FADH2<br>Folate; Tetrahydrofolate; 5,10-Methylenetetrahydrofolate<br>Thiamine diphosphate<br>Riboflavin<br>Biotin<br>Heme A<br>Vitamin B12 coenzyme<br>Undecaprenyl diphosphate<br>Ubiquinone-8<br>Pyridoxal phosphate |

| |
|---|
| Glutathionine reduced |
| Glutathionine oxidized |
| All-trans-Phytoene |
| Holo-carboxylase |
| $Co^{2+}$ (Cobalt) |
| $Mg^{+}$ (Magnesium) |
| $Ca^{2+}$ (Calcium) |
| $Mn^{2+}$ (Manganese) |
| $Fe^{3+}$ (Iron) |
| $Fe^{2+}$ (Iron) |
| $Zn^{2+}$ (Zinc) |
| $K^{+}$ (Potassium) |
| $Na^{+}$ (Sodium) |

[a] Composition based on the overall GC content of *S. meliloti* (1).

[b] The GC content for mRNA was estimated from the overall GC content of *S. meliloti* (1). The GC content of tRNA was estimated based on the GC content of the 10 most common codons in *S. meliloti* (1, 2). The GC content of rRNA was determined based on the *rrn* loci of *S. meliloti* (1). The overall composition of cellular RNA was determined assuming 80% rRNA, 15% tRNA, and 5% mRNA.

[c] The amino acid composition was estimated based on the codon usage of *S. meliloti* (2).

[d] The membrane lipid composition used was as previously determined for *S. meliloti* (3).

[e] A 4:1 ratio of low molecular weight (LMW) to high molecular weight (HMW) succinoglycan was set as previously determined (4).

[f] Each of these compounds were included at an equal, trace concentration in the biomass.

**Table S2.** Summary properties of the *S. meliloti* metabolic reconstruction iGD1348.

| | |
|---|---|
| **Genes** | 1348 |
| **Metabolites** | 1160 |
| Intra-cellular | 989 |
| Extra-cellular | 171 |
| **Reactions** | 1407 |
| Gene-associated reactions | 1164 |
| Metabolic reactions | 1019 |
| Gene-associated metabolic reactions | 982 |
| Transport reactions | 197 |
| Gene-associated transport reactions | 179 |
| Exchange reactions | 169 |
| Sink reactions | 8 |
| Source reactions | 2 |
| Biomass reactions | 1 |
| Other reactions | 11 |

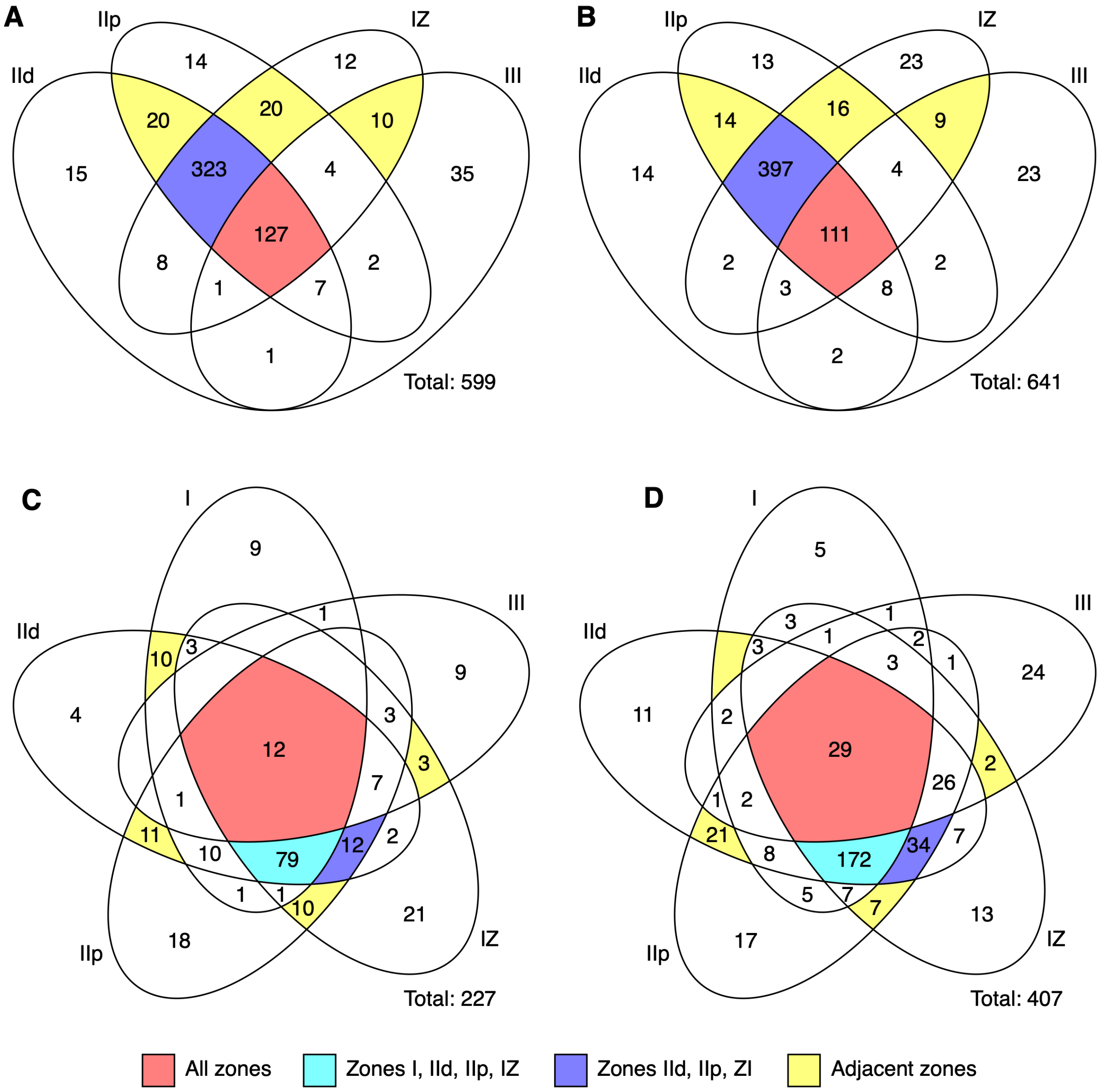


**Figure S1. Nodule zone-specific analysis of essential metabolism.** Venn diagrams are presented showing the overlap in genes or reactions predicted to be essential with FBA (growth rate ratio < 0.1 compared to the wild-type model) in each nodule zone. The total number of essential genes or reactions in each Venn diagram is indicated. Venn diagrams are shown for (**A**) *S. meliloti* genes, (**B**) *S. meliloti* reactions, (**C**) *M. truncatula* genes, (**D**) *M. truncatula* reactions. Notable sections of the Venn diagram are coloured according to the legend provided in the figure. The absence of a number in a Venn diagram section indicates there were zero genes or reactions with that characteristic.

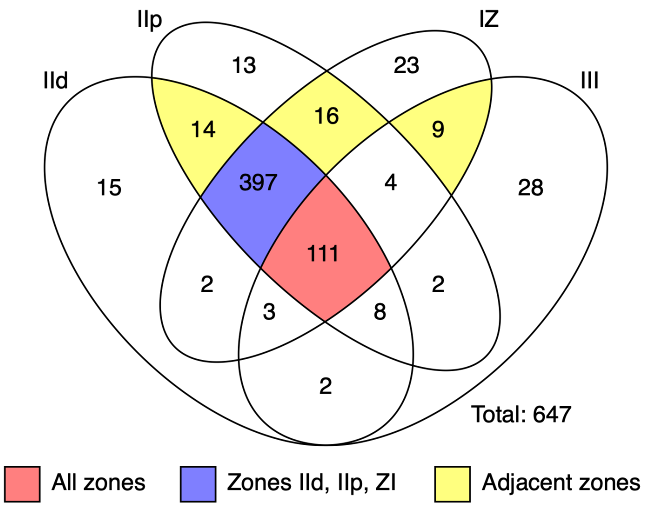


**Figure S2. Bacteroid robustness analysis summary.** An analysis was performed to evaluate the effects of perturbing the flux through each bacteroid reaction, specifically in each nodule zone, on the predicted rate of plant growth. A Venn diagram is presented to summarize the overlap in the "synergistic" bacteroid reactions, which are the bacteroid reactions that had to carry non-zero flux (i.e., they had to be active) in order for the predicted rate of plant growth to be at least 95% the maximal predicted plant growth rate. Notably, there is extremely high overlap between the essential reaction set (i.e., reactions whose removal results in a predicted plant growth rate less that 10% the maximal predicted plant growth rate) and the synergistic reaction set. Notable sections of the Venn diagram are coloured according to the legend provided in the figure.

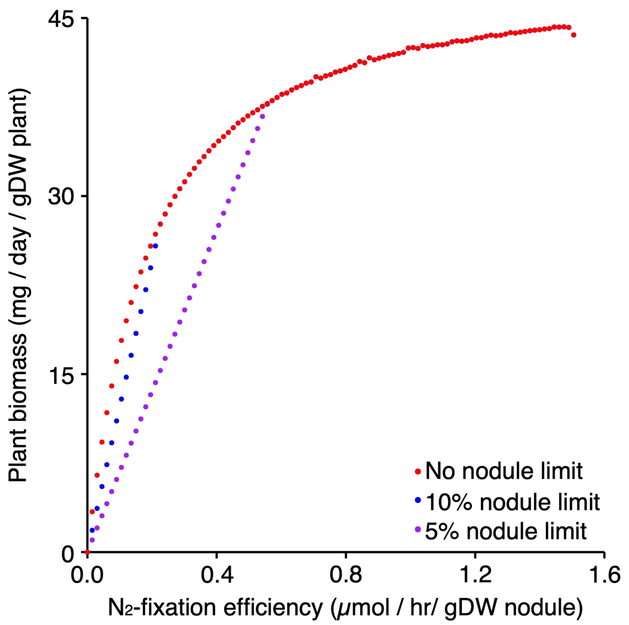


**Figure S3. Effect of $N_2$-fixation efficiency on plant biomass production.** The effect of $N_2$-fixation efficiency (rate of $N_2$-fixation per gram nodule) on the rate of plant growth, with the amount of nodule biomass optimized to maximize plant growth. A constant upper limit on the rate of oxygen uptake by zone III nodule tissue (adjusted based on the ratio between nodule and plant biomass) was used; see Figure 5D for a version without a limit on zone III oxygen uptake. Nodule biomass was either uncapped (red) or limited to 10% (blue) or 5% (purple) of the overall biomass.
