## Supplementary File S2 for "A Virtual Nodule Environment (ViNE) for modelling the inter-kingdom metabolic integration during symbiotic nitrogen fixation": README.txt

This archive contains the iGD1348 metabolic reconstruction as:
1) a SBML formatted file
2) an easy to read table in Excel format
3) a COBRA-formatted metabolic model saved as a MATLAB file.

The archive additionally contains the working version of the iGD1348 model, with unnecessary 'Unknown' GPRs removed from the model (i.e., GPRs such as [GeneA or Unknown] were replaced with [GeneA]), and with the gene 'NonEssential' removed.
