## Supplementary File S3 for "A Virtual Nodule Environment (ViNE) for modelling the inter-kingdom metabolic integration during symbiotic nitrogen fixation": README.txt

This archive contains two models. One of these is the working model used throughout this study (ViNE). The other is the full integrated model prior to constraining the nodule reaction space based on RNA-seq data (ViNE_precursor). The models are provided in two formats:
1) COBRA-formatted metabolic models saved as MATLAB files
2) SBML-formatted metabolic models, which were exported from MATLAB
